# *dnaE2* expression promotes genetic diversity and bacterial persistence in mycobacterial biofilms

**DOI:** 10.1101/2022.10.19.512518

**Authors:** S. Salini, Sinchana. G. Bhat, M. Nijisha, Reshma Ramachandran, Nikhil Nikolas, Krishna Kurthkoti

## Abstract

The complex cellular architecture and the microenvironments within the biofilm give rise to a population that is both physiologically and genetically heterogeneous. Transcriptome analysis of *Mycobacterium smegmatis* in biofilm culture and its transition phase into planktonic growth was performed to identify the genetic basis of heterogeneity in the biofilm. While there was an increase in the expression of mycobacterial mutasome consisting of *dnaE2, imuA*, and *imuB* in the biofilm, the mutation burden was less compared to the planktonic culture. Deletion of *dnaE2* causes lower mutation frequency and bacterial fitness in comparison to the parental strain in biofilm culture. The expression of *dnaE2* contributes to a slower bacterial growth rate, potentially promoting persister formation. Our study uncovers the multiple benefits of *dnaE2* expression in biofilm such as increasing genetic diversity and reducing growth rate; both of which are necessary for mycobacterial survival and adaptation.

## Introduction

Bacterial biofilms are complex, sessile bacterial communities with heterogenic populations that display different growth dynamics and physiochemical properties. The biofilm enables the bacterial community to not only withstand environmental stress such as nutritional starvation, host defence mechanism, antimicrobial agents but also propels them to evolve, and cause resurgence of an infection when favourable condition returns^1–3^. The biofilm formation is initiated by quorum sensing of autoinducer molecules such as acyl-homoserine lactone and nucleotide second messengers^4,5^ triggering the cell fate to change from a swarmer to sedentary life. This is followed by the production of an extracellular matrix consisting of polysaccharides, proteins and extracellular DNA (eDNA) that encapsulates the bacterial community as a single ecosystem^6–8^. The simultaneous occurrence of microenvironments, stochastic gene expression, and the emergence of mutants results in chemical and genetic heterogeneity within the bacterial population of biofilm ^9–13^. These differences in the biofilm confer bacteria the ability to respond to environmental stress and resist the action of antibiotics ^14^. Infections of pathogenic bacteria have become a serious public health concern for their ability to form biofilms both within the body and on medical devices^15,16^.

Members of mycobacteria form biofilms with a pellicle at the air–liquid interface in the absence of media compatible detergents such as Tween, or can be triggered by reductive stress^17,18^. While *in vitro* biofilms of mycobacteria have proved useful for understanding the bacterial physiology, its relevance in disease remained underappreciated till recently when Chakraborty et al., demonstrated the formation of biofilms of *M. tuberculosis* in infected lung tissues of mouse and humans^19^. During biofilm formation, the glycopeptidolipids facilitate mycobacterial attachment ^20–22^, followed by the synthesis of specialized short-chain C_56_-C_68_ free mycolic acids that are necessary for biofilm maturation^17^. Transcriptomic analysis of mycobacterial biofilms has shown the importance of the exochelin uptake system and GlnR-regulated genes in iron uptake, nitrogen assimilation, and resistance to peroxide stress ^23,24^. Biofilms from either fast or slow-growing mycobacteria contain populations of antibiotic-tolerant bacteria that are able to tolerate high levels of antibiotic treatment ^18,25,26^ posing a great concern to the outcome of drug treatment. In *M. tuberculosis*, thiol induced reductive stress accelerates biofilm formation that harbour drug-tolerant populations.

Multiple stress conditions, including reactive oxygen species within biofilms, are believed to the increase rate of mutagenesis, leading to genetic heterogeneity and evolution of bacteria within biofilms^3,27–30^. Although multiple studies have identified the genes required for biofilm formation and maturation, the basis of genetic heterogeneity and mutagenesis in mycobacterial biofilms is not well understood. In the present study, we investigated the role of *dnaE2* and its accessory factors *imuA* and *imuB* that constitutes the mycobacterial mutasome ^31,32^. Transcriptomic and reporter gene analyses showed the induction of the mutasome in the biofilm and its reduction during recovery into planktonic stage. Our findings further reveal that the expression of *dnaE2* provides a modest increase in resistance to rifampicin in wild-type biofilms compared to biofilms of Δ*dnaE2* strain. Interestingly, expression of *dnaE2* reduces bacterial growth rate, and its loss results in fitness defect in the mutant strain when competed with the parental strain in biofilm culture. Our study identifies the multifaceted roles of *dnaE2* in regulating mutagenesis and bacterial DNA replication in biofilm condition.

## Results

### Levels of ROS increases during *M. smegmatis* biofilm formation

We compared the gene expression of *M. smegmatis* during planktonic, biofilm and biofilm to planktonic transition (recovery stage) by performing RNA-Seq analysis on the isolated total RNA. We observed that the operon that encodes for oxidation of NADH (*nuoA*-*nuoM*) was upregulated and the gene cluster involved in ATP synthesis (*atpA-atpG*) was down-regulated in the 6^th^ day biofilm culture (Fig. 1A, and Supplementary Data sheet-1). Contrastingly, during the recovery stage when the biofilm was mechanically disrupted and subjected to planktonic growth, the gene expression of *nuoA-nuoM* gene cluster was down-regulated and that of *atpA-atpG* gene cluster was up-regulated (Fig. 1B and Supplementary Data sheet-2). We further verified this observation by determining the NADH levels at different stages of the biofilm using a mycobacterium optimized fluorescent NADH biosensor^33^. Microscopic analysis of the biofilm culture harbouring the biosensor revealed that there was a gradual increase in the NADH levels during biofilm formation (Fig. 1C, panels for 2^nd^ day and 4^th^ day). We also observed that there was a high degree of variation in the NADH level of 4^th^ day biofilm which could be arising as a consequence of the local heterogeneity in microenvironment. Additionally, the expression of genes involved in ROS detoxification such as, the *katG, sodC* and Mn dependent super oxide dismutase (*MSMEG_6427* and *MSMEG_6636*) were significantly downregulated in the biofilm culture (Fig. 1A). We reasoned that recycling of the NADH in the absence of ATP synthesis would be accompanied by the transfer of electrons to molecular oxygen resulting in generation of superoxide culminating in increase in ROS levels ^34,35^. To validate this observation, we disrupted the biofilm culture and stained the bacteria with ROS indicator CM-CH_2_DCFDA at different stages of biofilm maturation. We observed a gradual accumulation in ROS as the biofilm culture matured (Fig. 1D panels 2^nd^ day and 4^th^ day) favouring the predicted outcome from RNA-Seq and NADH biosensor data.

**Figure 1.**
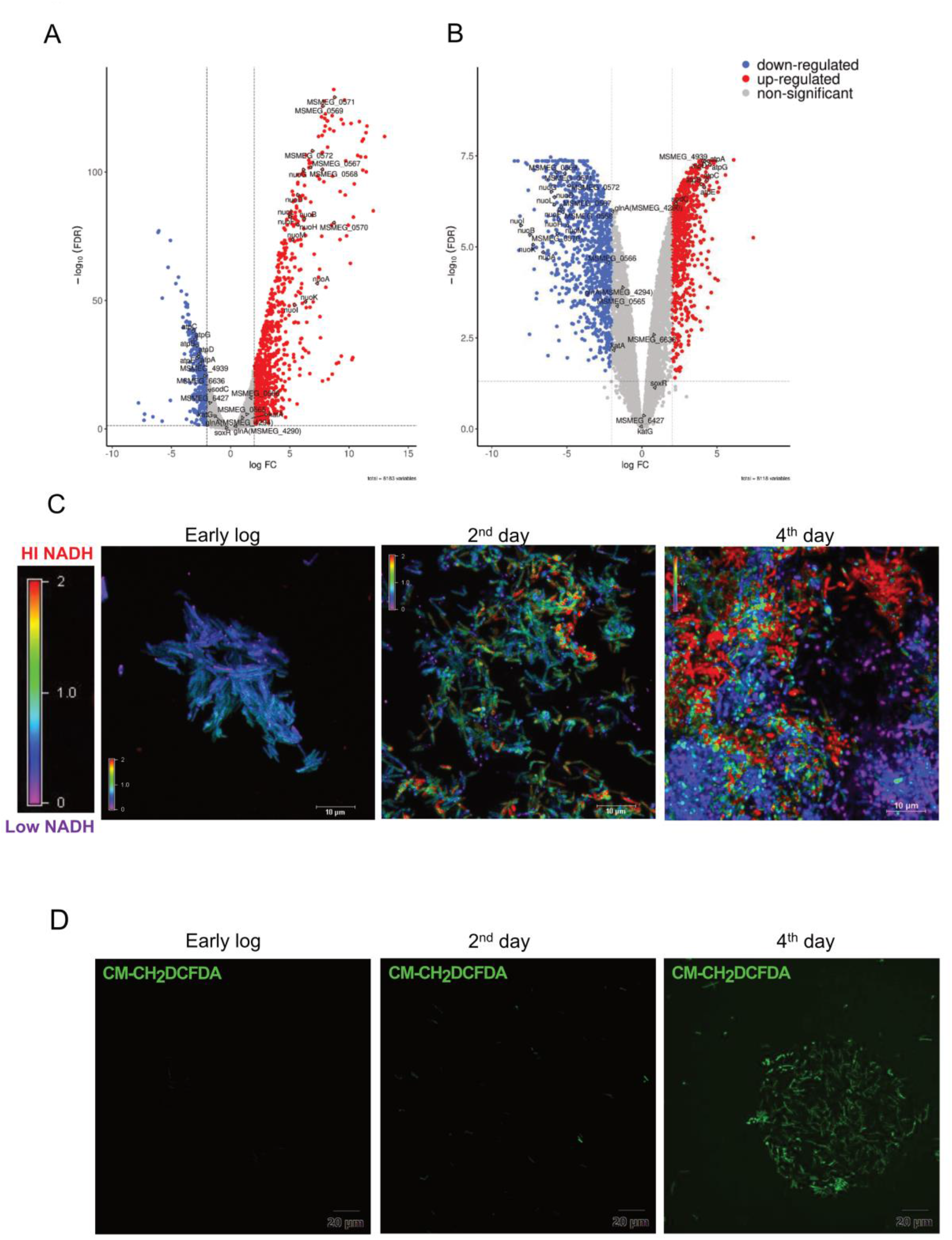
Determination of ROS level in biofilm using ROS indicator. A) Volcano plot depicting the fold up-regulated and down-regulated genes in biofilm culture in comparison to planktonic culture analyzed by RNA-Seq. B) Volcano plot of up-regulated and down-regulated genes during the recovery stage in comparison to biofilm culture. C) Intracellular redox status within the biofilm culture was determined using Peredox-NADH biosensor. *M. smegmatis* mc^2^155 was seeded for biofilm formation with a sterile grass cover slip in 24 well plate and the levels of NADH was monitored at 2^nd^ day and 4^th^ day intervals along with early-log phase culture as a control. The biofilm cultures were observed microscopically. The images were assigned a pseudo color for the representation of the NADH/NAD^+^ levels in individual cells and represented with the color heat map. The experiment was repeated with at least 2 replicates twice D) *M. smegmatis* mc^2^155 was seeded for biofilm in triplicates. After 2^nd^ and 4^th^ day of incubation, the cells were harvested, washed and stained with CM-CH_2_DCFDA ROS indicator and were subjected to CLSM. The fluorescent images were acquired and representative image is presented.

### DNA repair is down-regulated during biofilm formation but recovers when subjected to planktonic growth

Increased ROS levels result in DNA damage and induce DNA repair genes predominantly involved in the base excision repair pathway^36^. Therefore, we compared the expression profile of DNA repair genes in the total RNA of mature biofilm and the recovery stages by RNA-Seq analysis. To our surprise, genes involved in the GO repair pathway such as *mutY, mutM* and other DNA repair genes such as *ung, nei, xth* and *mfd* were down-regulated by 2-4 folds in the mature biofilms. However, during the recovery condition, the expression profile of the same DNA repair enzymes increased to the level of planktonic culture (Table-1). Interestingly, the error-prone polymerase *dnaE2* showed a reverse correlation i.e. the gene was induced during biofilm formation and downregulated during recovery phase. Along with *dnaE2*, its accessory factors *imuA* and *imuB* that constitute the mutasome cassette in mycobacteria were upregulated in biofilm (2-4 folds) but during the recovery phase, the expression was downregulated, approaching the level similar to that of planktonic culture (Table-1).

**Table 1.** List of genes involved in DNA repair that are differentially regulated in biofilm and recovery stages

| Gene name | Common name | Gene function | Planktonic to biofilm | Biofilm to Recovery | Planktonic to Recovery |
| --- | --- | --- | --- | --- | --- |
| *MSMEG_2399* | *ung* | uracil-DNA glycosylase | 0.61 | 1.90 | 1.04 |
| *MSMEG_2419* | *mutM* | formamidopyrimidine-DNA glycosylase | 0.77 | 1.96 | 1.34 |
| *MSMEG_4683* | *nei1* | putative formamidopyrimidine-DNA glycosylase | 0.30 | 7.62 | 2.09 |
| *MSMEG_6187* | *nth* | endonuclease III | 0.60 | 2.42 | 1.29 |
| *MSMEG_0829* | *xth* | exodeoxyribonuclease III | 0.33 | 3.96 | 1.19 |
| *MSMEG_2765* | *dut* | dUTP hydrolase | 0.49 | 3.96 | 1.74 |
| *MSMEG_5148* | *mutT2* | CTP pyrophosphohydrolase | 0.26 | 2.55 | 0.66 |
| *MSMEG_0790* | *mutT3* | hydrolase, NUDIX family protein | 0.56 | 1.62 | 0.81 |
| *MSMEG_6927* | *mutT4* | MutT/nudix family protein | 0.53 | 2.21 | 1.04 |
| *MSMEG_3078* | *uvrC* | excinuclease ABC, C subunit | 0.68 | 2.40 | 1.45 |
| *MSMEG_5423* | *mfd* | transcription-repair coupling factor | 0.25 | 11.04 | 2.45 |
| *MSMEG_3172* | *dinX(dinB1)* | DNA polymerase IV 1 | 0.58 | 4.64 | 2.38 |
| *MSMEG_1633* | *dnaE2* | DNA polymerase III, alpha subunit, putative | 2.12 | 0.75 | 1.44 |
| *MSMEG_1620* | *imuA'* |  | 5.03 | 0.29 | 1.35 |
| *MSMEG_1622* | *imuB* | putative DNA repair polymerase | 2.74 | 0.57 | 1.39 |

### The mycobacterial error-prone polymerase *dnaE2* and its associated proteins are induced during biofilm formation

The expression of *dnaE2* constitutes the SOS response which is dependent on *recA*. To analyse the *recA* expression during different stages of biofilm, we generated a destabilized mClover reporter fused to *M. smegmatis recA* promoter sequence (*P*_*recA*_). The *M. smegmatis* strain harboring the *P*_*recA*_∼mClover reporter plasmid was seeded for biofilm formation and the reporter signals were collected after every 2 days. We observed high expression of *recA* in the early stages of biofilm (Fig. 2A panel 2^nd^ day) that progressively decreased as the biofilm matured (Fig. 2A panels 4^th^ day and 8^th^ day). The expression of DnaE2 along with the other accessory factors ImuA’ (MSMEG_1620) and ImuB (MSMEG_1622), results in mutagenesis during the SOS response in mycobacteria ^31,32^. We therefore checked for the expression of *dnaE2* and *imuA* at different stages of biofilm formation (Fig. 2B and 2C respectively) using gene-specific promoters fused to destabilized version of mCherry. Microscopic analysis of the biofilm revealed that the expression of *dnaE2* was minimal at 2^nd^ day and subsequently increased with the maturation of the biofilm (Fig. 2B panels 4^th^ day and 8^th^ day). The expression pattern of *imuA* more or less overlapped with the expression pattern of *dnaE2* with consistent higher levels after 4^th^ day in biofilm and increasing further at 6^th^ day (Fig. 2C). The *imuA* and *imuB* constitute an operon and therefore the expression pattern of *imuB* could not be determined by reporter analysis and we believe that the expression of *imuB* would be similar to that of *imuA*. The reporter expression revealed that there was a delay in the expression of *recA* and its downstream target *dnaE2* (2^nd^ day for *recA* and 4^th^ day for *dnaE2* in Fig. 2A and Fig. 2B respectively). To validate if the observed temporal difference in the expression of *recA* and *dnaE2* was not due to experimental variation arising in two different strains, we generated a dual reporter strain harboring *P*_*recA*_∼mClover and *P*_*dnaE2*_∼mCherry and observed the expression pattern of the individual genes. In agreement with our individual reporter findings, the expression of *P*_*recA*_∼mClover preceded the expression of *P*_*dnaE2*_∼mCherry (Fig. 2D panels 2^nd^ to 8^th^ day). Also, deletion of *recA* resulted in significant reduction of *dnaE2* expression in biofilm in agreement with earlier reports ^31^ (Fig. S1 panels 2^nd^ day, 4^th^ day and 8^th^ day). Furthermore, the expression of the cell division checkpoint regulator of mycobacteria (*MSMEG_3644*) that functions as a transcriptional activator of *dnaE2* was also induced by ∼5 folds during biofilm formation (Table-2) ^37^.

**Table 2.**
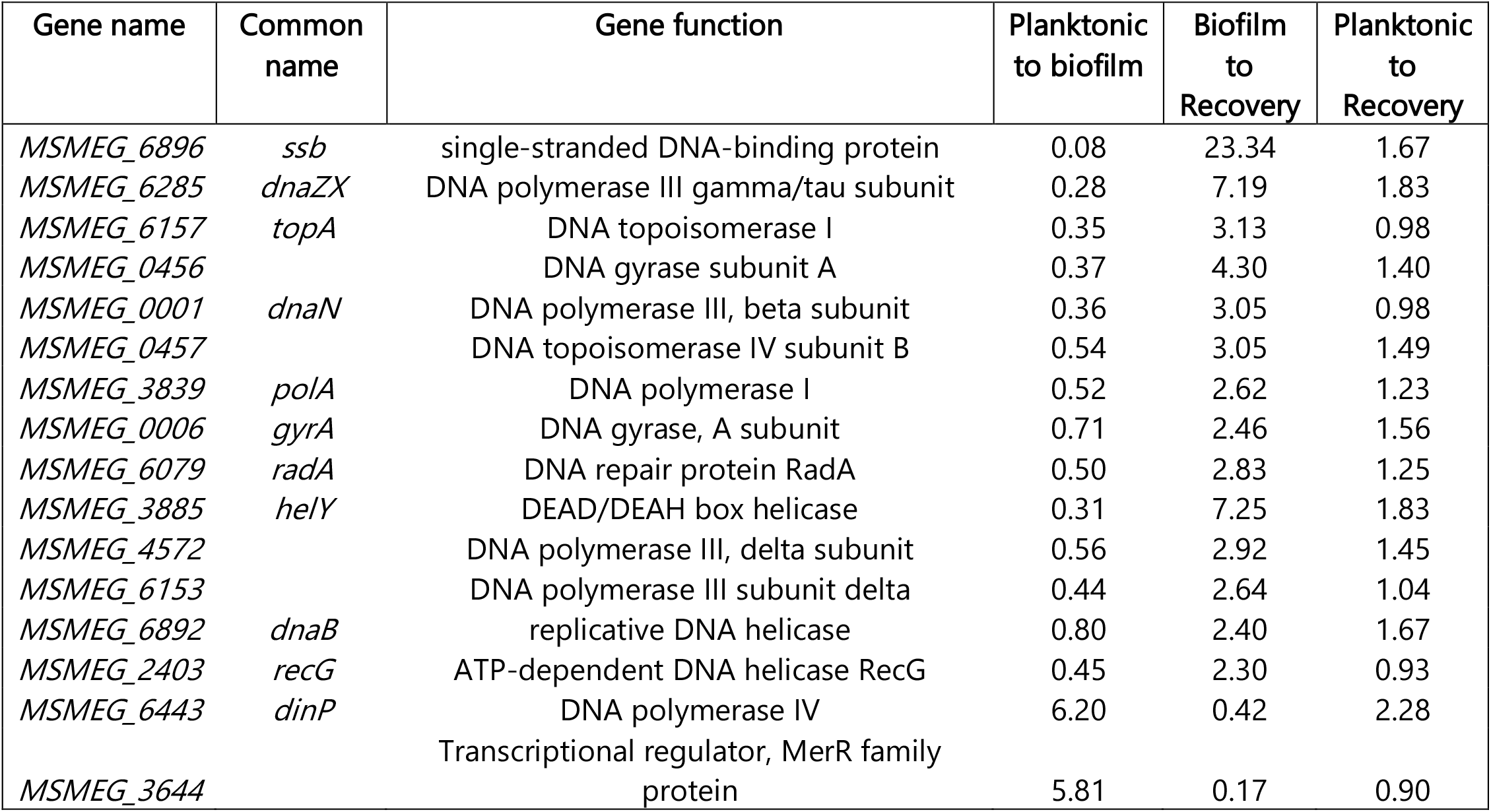
List of genes involved in DNA replication that were differentially regulated in biofilm and recovery

**Figure 2.**
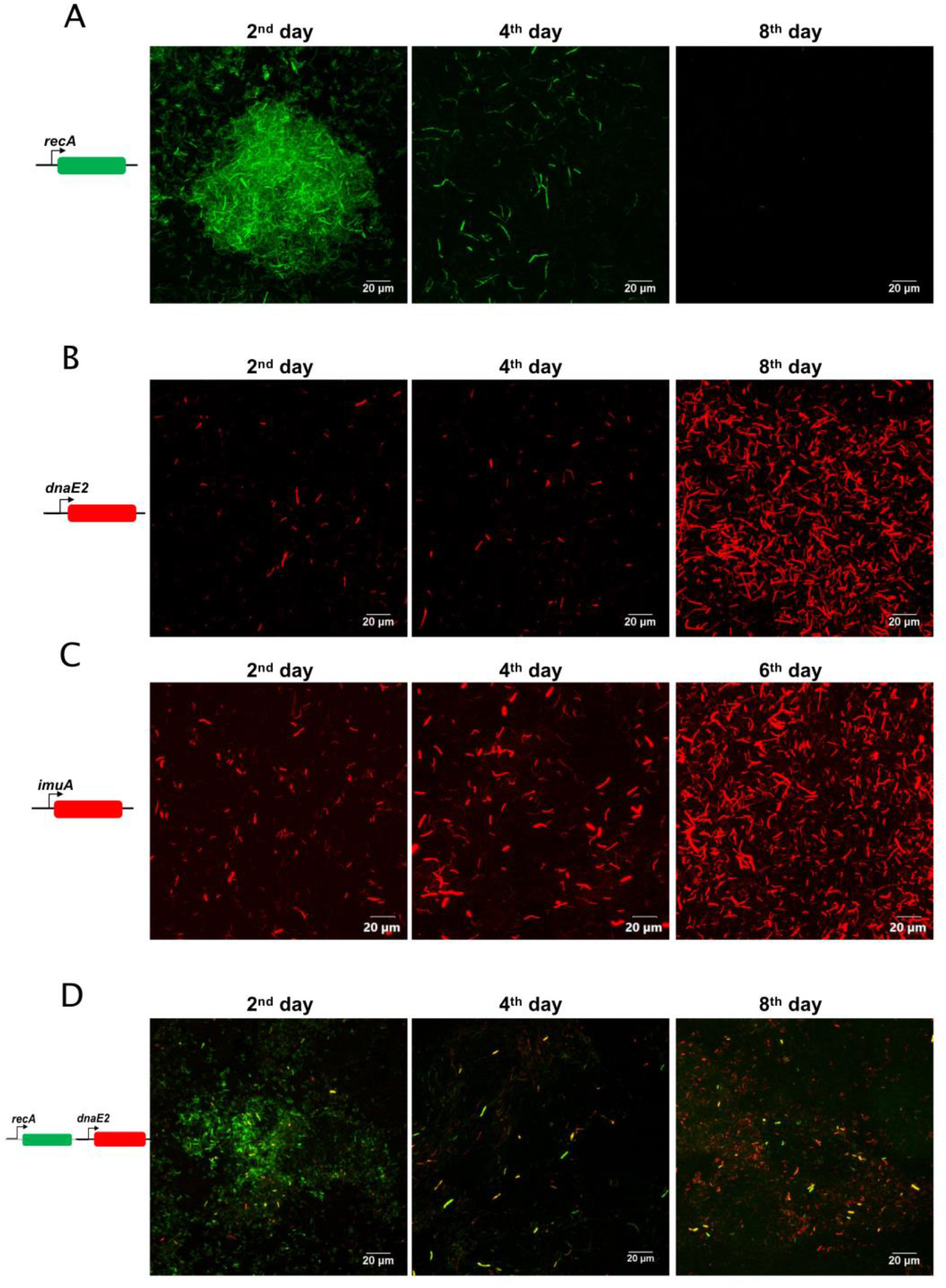
Expression analysis of *recA, dnaE2* and *imuA* in biofilm. A) *M. smegmatis* harboring pMV262 *P*_*recA*_*∼*mClover was seeded for biofilm with a sterile glass coverslip in a 24 well plate and incubated for 8 days. After incubation, the media was removed and the biofilm layered on the cover slip was imaged. The fluorescent images were acquired at defined time points and a representative image is presented. B) *M. smegmatis* with pMV262 *P*_*dnaE2*_*∼*mCherry was seeded for biofilm as mentioned before and fluorescent images were acquired at defined time points and representative image presented C) *M. smegmatis* with pMV262 *P*_*imuA*_*∼*mCherry was seeded for biofilm formation for 6 days and fluorescent images were acquired at defined time points and representative image is presented. D) *M. smegmatis* harboring dual reporter pMV262 *P*_*recA*_*∼*mClover and *P*_*dnaE2*_*∼*mCherry was seeded for biofilm and incubated for 8 days. The fluorescent images corresponding to the expression of *P*_*recA*_ (green) and *P*_*dnaE2*_ (red) were acquired at defined time intervals and the merged image is presented.

### Contribution of DnaE2 expression to bacterial heterogeneity in biofilm

Transcriptome analysis from biofilm condition revealed that the expression of DNA repair enzymes was down-regulated while that of the mycobacterial mutasome consisting of *dnaE2, imuA* and *imuB* was increased (Table-1). This observation made us to hypothesize that the two can synergistically promote mutagenesis in biofilm cultures. Accordingly, we determined the occurrence of spontaneous mutants to antibiotics as a measure of mutagenesis. Surprisingly, the mutation frequency of rifampicin resistance (Rif^R^) between planktonic and biofilm stages was insignificant in WT strain (Fig. 3A). We repeated the experiment and scored for mutations that confer resistance to ciprofloxacin (Cip^R^) and observed that the mutations in planktonic culture was considerably higher than the biofilm culture (Fig. 3B). We deliberated that the observed lower mutation frequency in biofilm cultures could arise from the death of bacteria within the biofilms that harbour Rif^R^ or Cip^R^ mutants. To determine the extent of dead bacteria in the biofilms, we stained the 6 day biofilm culture with a combination of Syto9 and Propidium iodide dyes (Live/Dead staining) and performed microscopic analysis. We observed that there was significant population of dead bacteria in the biofilm culture compared to the planktonic culture (Fig. 3C, right panel). Next, we monitored the contribution of *dnaE2* to mutagenesis in biofilm by determining the mutation frequency of WT, *ΔdnaE2* and *ΔdnaE2*::*pdnaE2* (complemented) biofilm cultures to Rif^R^ and Cip^R^. While the difference in the mutation frequency of WT and *ΔdnaE2* in biofilm condition to Rif^R^ was quite evident (Fig. 3D; median value of 1.28×10^−8^ for WT in comparison to 4.9×10^−9^ for *ΔdnaE2*) the observed difference in mutation frequency to Cip^R^ was less apparent (Fig. 3E, medial value of 8.9×10^−8^ for WT in comparison to 5.8×10^−8^ for *ΔdnaE2*). Since the expression of *dnaE2* reduces during recovery into planktonic phase of growth (Table-1), we performed a time course expression analysis of *dnaE2* during recovery stage using a dual reporter harboring a *P*_*hsp*_∼mClover and *P*_*dnaE2*_∼mCherry^38^. While there was expression of *P*_*dnaE2*_∼mCherry at the time of biofilm disruption (0 h) the expression significantly reduced between 3 to 6 hrs (Fig. 3F, panels 0 hr-6 hr). We believe that the induction of DNA repair and downregulation of mutasome in recovery stage could also contribute to reduction in the mutation frequency. To account for the potential loss of Rif^R^ and Cip^R^ phenotype due to DNA repair during processing and plating in the mutation frequency experiment, we attempted to score for “gain-of” fluorescence mutation in the biofilm using a non-fluorescent version of GFP (GFP*). Transformants of *M. smegmatis* mc^2^ 155 and *ΔdnaE2* harboring GFP* were subjected to biofilm formation and observed for appearance of green fluorescence. While the reversion rate of GFP* was very low even in WT, we could not observe any GFP reversion in the mutant strain (Fig. S2, panels of WT and *ΔdnaE2* respectively). Based on these findings we conclude that there is a minor contribution of *dnaE2* mutasome towards generation of genetic heterogeneity in the biofilm.

**Figure 3.**
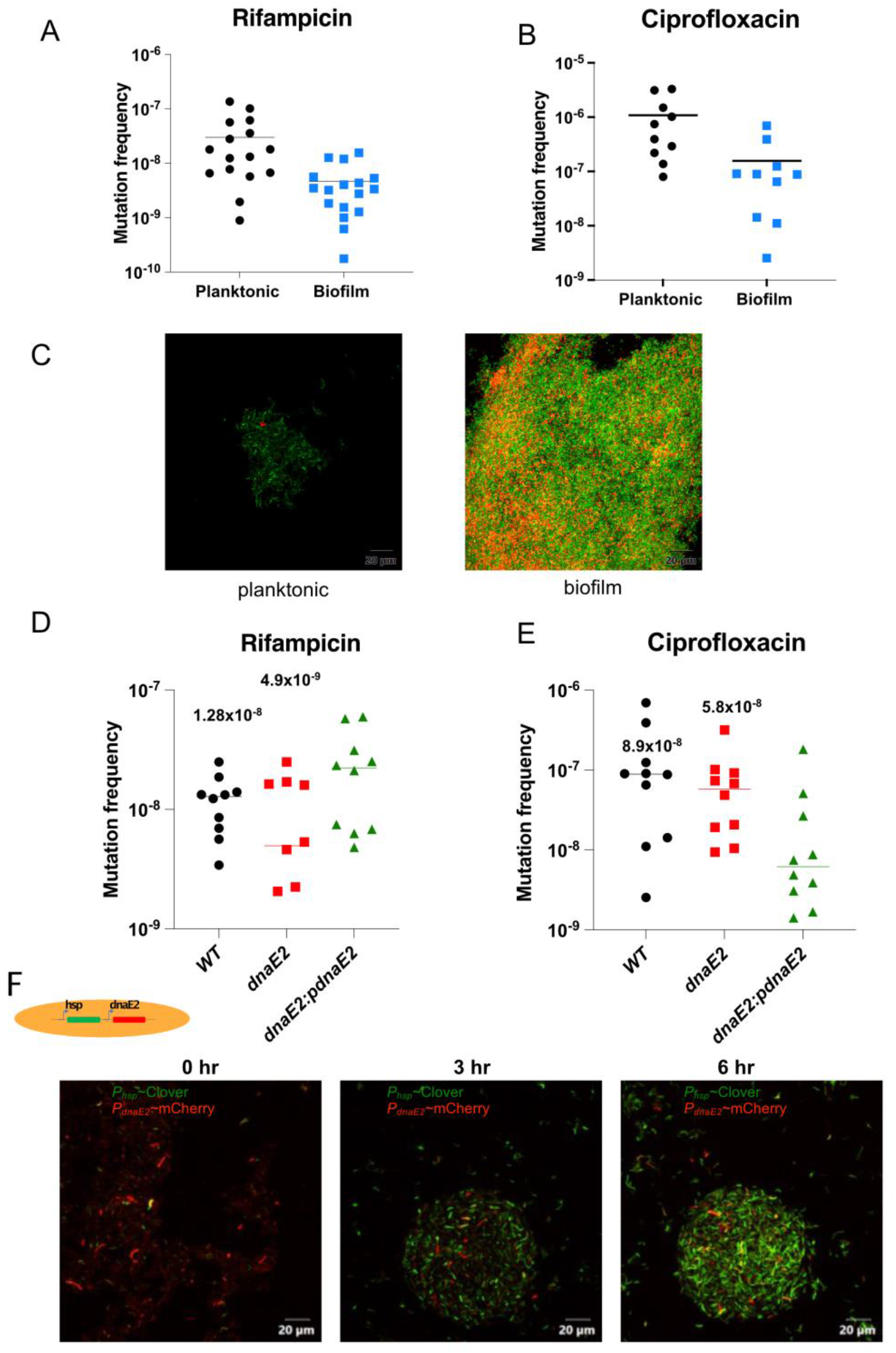
Contribution of *dnaE2* expression to bacterial heterogeneity in biofilm. A) Ten or more replicates of each strain were subjected to biofilm formation and plated on to either Cip or Rif plates. The mutation frequency was determined by the ratio of number of mutants to the viable count for each replicate. The individual values were plotted on a scatter plot using GraphPadPrism^®^ software v8 and the corresponding median value for each set indicated above the plot. (*P values*, * *P*<0.05). A) Mutation frequency of *M. smegmatis* (wild-type) for Rif. B) Mutation frequency of *M. smegmatis WT* for Cip. C) Three replicates each of mid-log phase planktonic and 6d biofilm cultures were harvested and stained with Syto9/PI dyes (Live/Dead staining) and subjected to confocal microscopy to visualize live (green) and dead (red) bacterial population. The experiment was repeated at least twice and representative images are shown D) Mutation frequency of *M. smegmatis WT, ΔdnaE2 and ΔdnaE2*::*pdnaE2* for Rif. E) Mutation frequency of *M. smegmatis WT, ΔdnaE2 and ΔdnaE2*::*pdnaE2* for Cip. E) Time course expression analysis of *dnaE2* during biofilm recovery stage of *M. smegmatis* using *P*_*hsp*_∼mClover and *P*_*dnaE2*_∼mCherry dual reporter in 7H9T medium.

### *dnaE2* is required for bacterial fitness in biofilm culture

To understand the importance of *dnaE2* expression during biofilm formation, we determined the fitness of *dnaE2* mutant strain in comparison to WT by co-culturing them within the biofilm. To perform this competition study, we generated fluorescently labelled strains of WT and Δ*dnaE2* encoding mClover and mCherry respectively through the *P*_*hsp*_ promoter. The culture containing equal proportion of WT and Δ*dnaE2* were allowed to form biofilm and the population density was observed microscopically. During the initial stages, the bacterial population density was ∼50% for the WT and the Δ*dnaE2* strains (Fig. 4A, panel i and Fig 4C, 0 passage). However, upon repeated passaging, the density of Δ*dnaE2* (red bacteria) drastically diminished to ∼5% by fifth passage in biofilm (Fig. 4A panel ii-iii and Fig. 4C, 5^th^ passage). To rule out the potential impact of mClover or mCherry on the fitness of bacteria, we competed WT strain harboring mClover and mCherry and did not observe significant reduction in the population density at 5^th^ passage (Fig. 4B panel i-iii). We checked the growth kinetics of WT, Δ*dnaE2* and complemented strains in liquid medium to determine if the mutant strain exhibited any growth defects. There was no significant reduction in growth rate of the Δ*dnaE2* strain when cultured independently in liquid medium (Fig. 4D). These findings ruled out that the observed fitness defect of the Δ*dnaE2* in the biofilm when competed against WT is not due to its inherent reduced growth defect.

**Figure 4.**
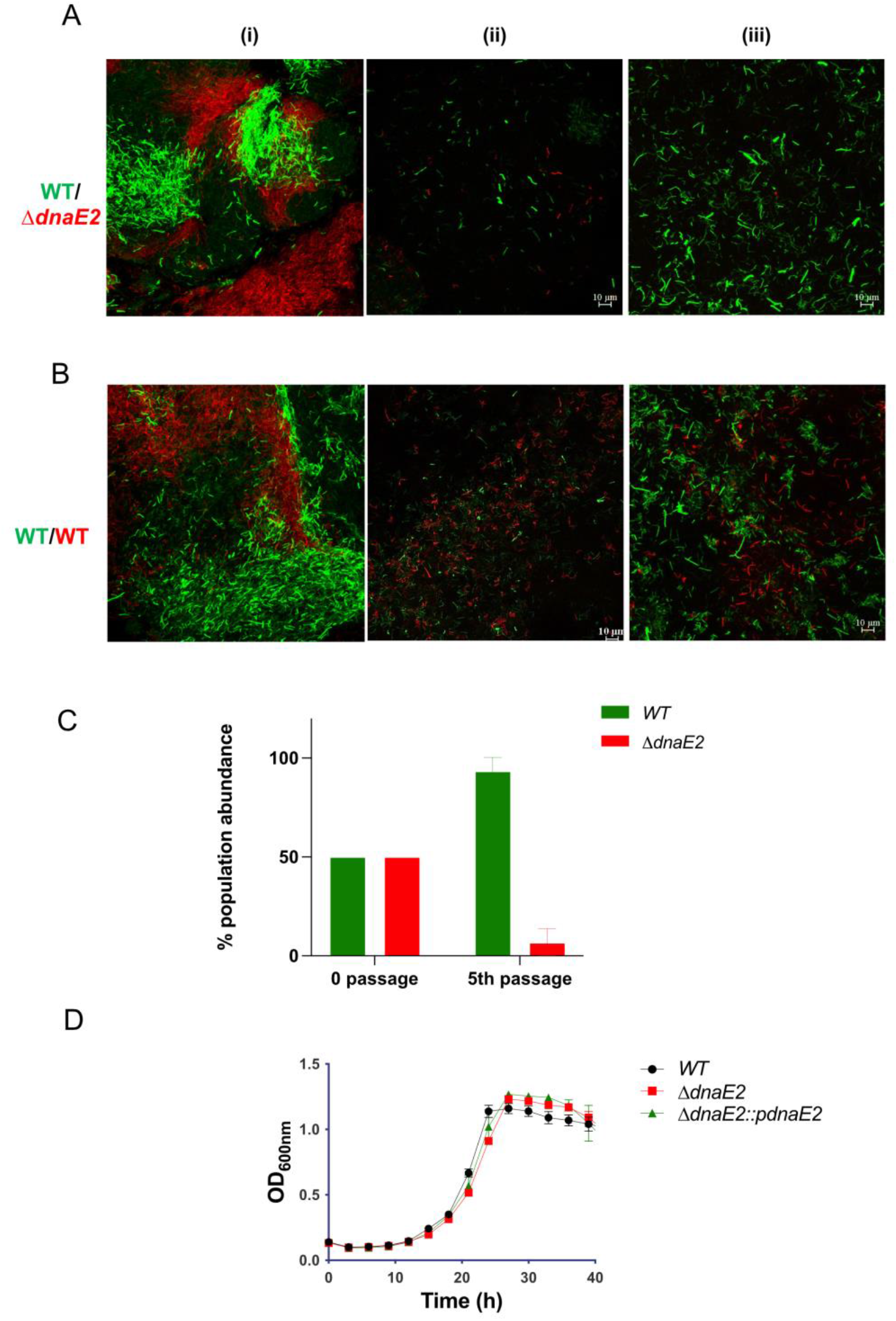
Competition analysis of WT persister with Δ*dnaE2* biofilm population. A-B) *M. smegmatis* wild-type and Δ*dnaE2* strains expressing short half-life variants of either mClover and mCherry respectively through *P*_*hsp*_ were mixed in equal proportions and co-cultured continuously for five passages. Microscopic images of the mixed population at 1^st^ 3^rd^ and 5^th^ passages were acquired. The experiment was performed at least twice C) The population abundance of *WT* and Δ*dnaE2* strain at the beginning and at 5^th^ passage was plotted from three replicates. Representative graph from two independent experiments are presented. The experiment was repeated twice with two replicates. D) Growth profile of *M. smegmatis* strains in 7H9T medium. Five replicates of each strain was diluted to an final OD_600_ ∼0.02 and seeded to individual cells of Honeywell comb plate. The plate was incubated at 37 °C with constant shaking and the increase in growth was monitored by absorbance measurement at 600 nm in Bioscreen C instrument (Bioscreen, Talekatu Finland).

### Expression of *dnaE2* results in reduced bacterial growth rate

The mRNA levels of genes involved in replication such as *ssb, topA, gyrA, dnaN* were significantly downregulated while the expression of *MSMEG_3644* that inhibits bacterial division was induced in biofilm condition (Table-2). The induction of error-prone polymerase has been shown to reduce the rate of DNA replication in *E. coli* ^39^. Since the expression of *dnaE2* was observed as early as 2 days, we reasoned that its expression and binding to the genomic DNA could retard the replication and reprogram the bacteria for slower growth rate. We compared the growth of mycobacteria in planktonic and biofilm culture by both turbidity and dilution plating methods. Bacterial cultures for planktonic and biofilm cultures were set at 0.02 OD (0 day) and the viability was ∼ 5×10^6^ CFU/mL. While the planktonic culture reached saturation by 2^nd^ day (∼2×10^9^ CFU/mL) followed by death phase by 6^th^ day, the biofilm culture reached late log-phase by 4^th^ day (Fig. 5A) with lesser viability in comparison to its optical density (3.3×10^7^ CFU/mL) implying the culture had substantial dead bacteria and the matrix components contributing to the apparent high optical density of the culture. Comparing the viable counts of planktonic and biofilm cultures also showed that the increase in viable counts in biofilm culture was significantly slower compared to the planktonic culture (Fig. 5B, red bars to green bars). To rule out the differences in the culture conditions of biofilm and planktonic stages for the observed slow growth rate, we treated planktonic culture of WT strain to Cip for 48 h that generates a persister stage with robust expression of *dnaE2*. Recovery of antibiotic persister (AP) cultures of WT in antibiotic free medium was significantly slower and took nearly 6 days for attaining stationary phase compared to 4 days for similarly treated and recovered *ΔdnaE2* strain (Fig. 5C and 5D). Although this variation could arise due to differences in the number of surviving bacteria upon antibiotic’s lethal action, we considered the possibility that a reasonable proportion of APs entered a “differentially detectable” state (DDS) or viable but nonculturable state (VBNC). It has been demonstrated that bacteria in DDS display high ROS ^40^ and co-exist with antibiotic persisters ^41^. We therefore analyzed the viable status of APs by live/dead staining in the Cip treatment and observed that the proportion of live bacteria (green stained) was comparable between the wild-type and *ΔdnaE2* strains (Fig. 5E, panel Cip and its corresponding zoomed area). These observations indicate that *dnaE2* expression constitutes an additional factor in the formation of slow growing persister population within the biofilm.

**Figure 5.**
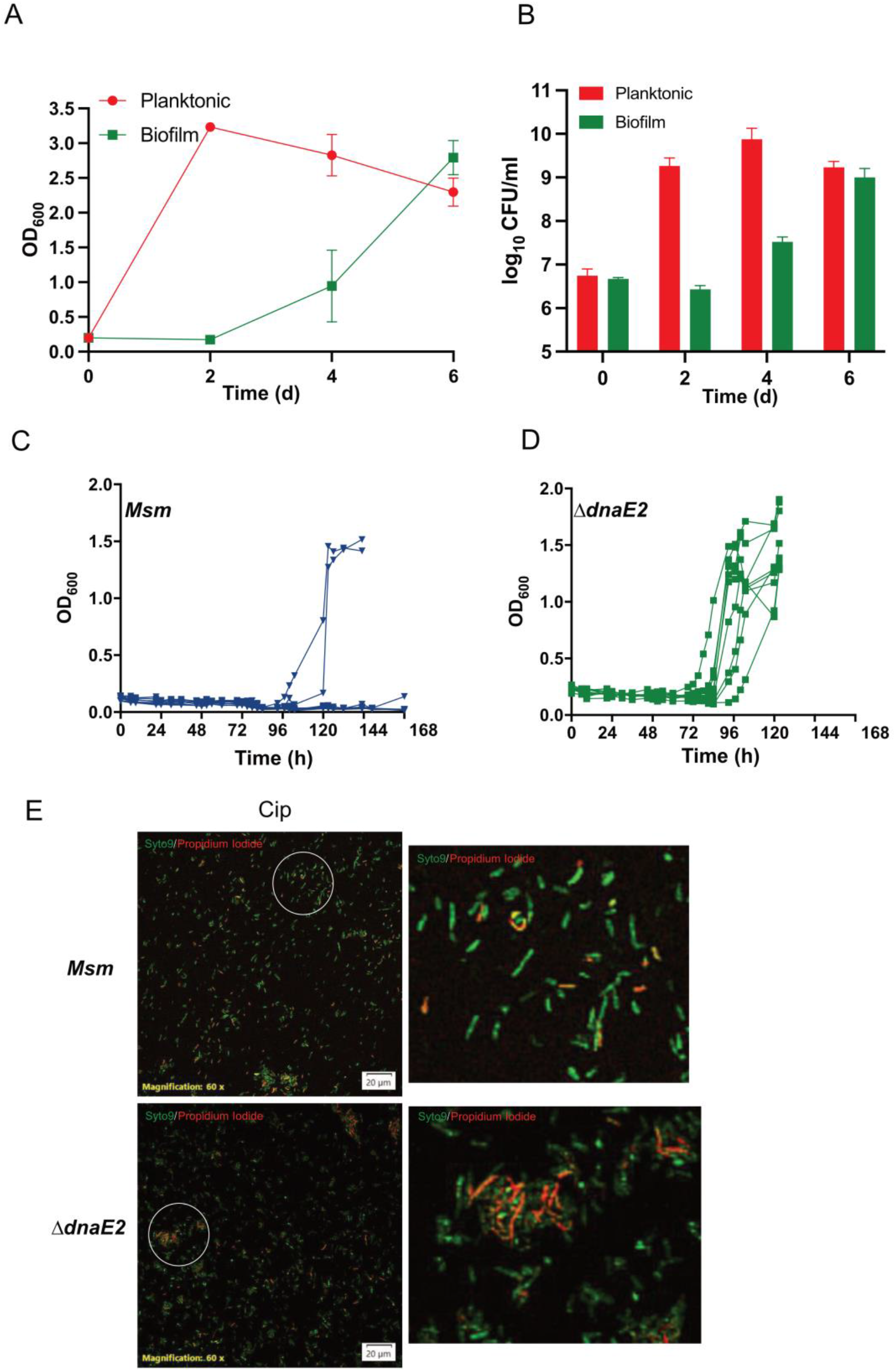
Negative impact of *dnaE2* expression on mycobacterial growth. A) Growth curve analysis of planktonic and biofilm culture. OD_600_ of the cultures were measured on 2^nd^, 4^th^, and 6^th^ day. The growth curve for each replicate was plotted on a scatter plot using GraphPadPrism^®^ software and the corresponding median value for each set indicated above the plot. (*P values*, * *P*<0.05). B) the viable count (CFU/mL) of planktonic and biofilm culture. The individual value of log CFU/ml was plotted against the time points in GraphPadPrism^®^ software v8 (*P values*, * *P*<0.05). C and D) Growth curve analysis of Cip persisters during recovery phase. Antibiotic persister cells were generated for WT (C) and *ΔdnaE2* (D) by Cip treatment for 48 h and recovered in plain medium. The OD_600_ of the cultures was measured at regular intervals and the growth curve for each replicate was plotted. The images presented is a representative of two independent experiments. E) From an independent experiment, the Cip persisters of *M. smegmatis* wild-type, *ΔdnaE2* were subjected to Live/Dead staining. The experiment was repeated at least twice with 3 independent replicates. A zoomed image of *M. smegmatis* wild-type and *ΔdnaE2* Cip persisters are shown from the circled region.

## Discussion

The intricate 3D architecture of biofilm results in generating microenvironments due to which bacteria display both chemical and genetic heterogeneity ^9,13,14^. In the present study, we investigated the involvement of mycobacterial error-prone polymerase in causing genetic mutations in biofilm. Our findings show that during biofilm formation the levels of ROS increases, inducing SOS response, and resulting in expression of DnaE2. The reduced DNA repair and increased levels of DnaE2 can synergistically promote mutagenesis in the biofilm. Additionally, the occupation of DnaE2 on the genomic DNA may conflict with the function of replication polymerase DnaE1 resulting in reduced bacterial growth. However, when the biofilm bacteria are subjected to planktonic growth, the DNA repair enzymes are induced along with proteins involved in replication with the concomitant reduction of DnaE2 promoting bacterial growth (Fig. 6).

**Figure 6.**
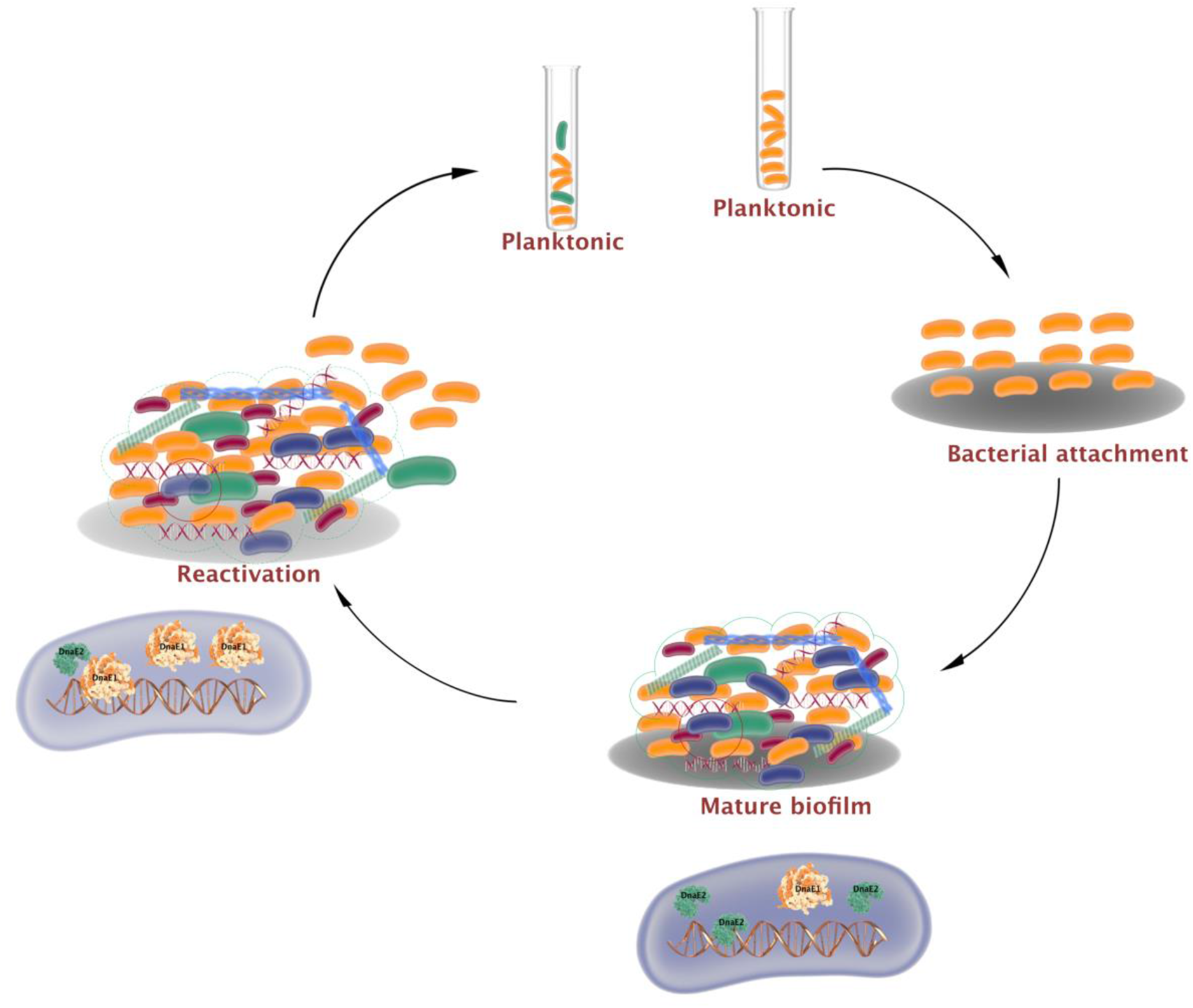
Mutagenesis and bacterial growth arrest by *dnaE2* expression in biofilm. Biofilm formation initiates with the attachment of mycobacteria to a substratum. During maturation, an extracellular matrix encloses the heterogenous population with dead bacteria (red), growth-arrested bacteria (blue), and mutant bacteria (green) generated by reduced DNA repair and induction of error-prone polymerase DnaE2. The growth arrest within the sub-population emerges from the conflict between DnaE2 and the house keeping DnaE1. During reactivation of biofilm to planktonic growth, expression of DnaE2 is repressed, and balance of polymerases shifts in favour of DnaE1 resulting in active multiplication.

Many bacteria form biofilms in response to stress such as antibiotics and oxidative agents that induces SOS response ^42–44^. To further understand the role of SOS response and its contribution to mutagenesis, RNA-Seq analysis was performed on biofilm or recovered culture of *M. smegmatis* and we observed induction of *recA* and the mycobacterial mutasome during biofilm formation (Fig. 2 and Table-1). We were intrigued by the induction of SOS regulon genes in the biofilm since there was no additional stress administered to the culture. Gene expression analysis showed increased expression of NADH oxidase enzymes with concomitant decrease in ATP synthesis resulting in increased ROS levels during biofilm formation (Fig. 1A and 1C). ROS has been speculated to function as a signalling molecule to induce biofilm formation^45,46^ and in *M. avium* presence of auto inducer molecules can increase ROS and stimulate biofilm formation ^47^. To overcome peroxide stress generated within biofilms, Yang et al., have shown the expression of Gln dependent regulon (*MSMEG_ 0565-0572*) during biofilms confers resistance to peroxide and helps in nitrogen assimilation ^24^. In agreement with their observation, our RNA-Seq analysis revealed that the same genes were highly upregulated during biofilm formation (Fig. 1A). The presence of ROS can also contribute to the stability of *Streptococcus spp* biofilm by stimulating the release of eDNA ^48,49^. The potential role of ROS in release of eDNA in mycobacteria has not been explored so far but it is known to inflict DNA damage including strand breaks and oxidation of DNA bases.

Mycobacterial genomes being GC rich are highly sensitive to oxidation of Guanine resulting in formation of 8-oxo-Guanine residues that can interfere with replication. To overcome the DNA damages an induction of DNA repair response was expected during biofilm. However, DNA repair enzymes belonging to base excision repair pathway were downregulated but the error-prone polymerase *dnaE2* and its accessory factors were upregulated (Table-1). Interestingly, the expression of *recA* which is the master regulator of SOS response and necessary for expression of *dnaE2* was significantly down-regulated at 4^th^ day and subsequent time points (Fig. 2A and 2D panels for 2^nd^ day and 4^th^ day and Table-1). This observation is in agreement with Yang et al 2017, where a time course RNA expression profile in developing biofilms showed that RecA was significantly induced in early timepoints of biofilm culture (∼ 8 folds) but dropped to ∼0.7 fold at later stages ^22^. On the contrary, we observed that the expression of *dnaE2* steadily increased from 4^th^ day to 8^th^ day which is inversely related to *recA*’s expression. To understand the rationale behind this inconsistency we looked for potential regulators that could induce *dnaE2*. A recent study has shown that over-expression of a MerR family transcription regulator Rv1830 (MSMEG_3644) acts as a mycobacterial cell cycle checkpoint by preventing mycobacterial cell division and simultaneously induces expression of *dnaE2* and *imuB* ^37^. Our RNA-Seq data revealed that the expression of *MSMEG_3644* was induced by ∼5 folds in the biofilm stage but was repressed during the recovery stage (Table-2).

The oxidative damage within the biofilm has been shown to cause increased mutagenesis in biofilm communities of *Pseudomonas* and *Staphylococcus* ^29,50^. Further the mutations within the biofilm could result in *de novo* emergence of drug resistance and other stress tolerating variants that would have a competitive advantage ^3,27,51^. Since mutagenesis within mycobacterial biofilms has been relatively underexplored and that the mutasome of mycobacteria was induced, we determined the mutation frequency by scoring for spontaneous resistant mutants to Rif and Cip in the biofilm cultures. Surprisingly, the mutation frequency was significantly lesser than the planktonic culture (Fig. 3A and 3B) in spite of having higher expression of *dnaE2* and *imuA* (Fig. 2B and 2C respectively). There are conflicting reports about bacterial mutagenesis in biofilms. While Conibear *et al*., have reported ∼100 fold increase in mutation in microcolony foci^30^, Garcia-Castillo *et al*., have observed reduced mutagenesis of biofilm culture compared to planktonic cultures in clinical isolates of *Pseudomonas* reflecting our results ^52^. Multiple factors could lead to such an outcome. In the antibiotic-based mutation scoring assay, only mutations that confer resistance produce colonies and often resistance to Rif or Cip generate mutants with high fitness cost ^53,54^ and may be competed out during the biofilm formation stage. Additionally, death of antibiotic-resistant bacterium in the biofilm would be reflected as an apparent decrease in mutation frequency (Fig. 3C). Recent studies have shown that bacteria released from biofilm are physiologically distinct and are more sensitive to antibiotics than their planktonic counterparts^55^ and this feature could further complicate the interpretation of antibiotic resistance based read-out. RNA-Seq and time course gene expression analysis of (Table-1 and Fig. 3F) suggests a further reduction of mutations due to increased DNA repair and reduction in *dnaE2* expression. Using the fluorescence reversion assay, we could visualize the appearance of GFP positive cells only in the WT strain but not in the mutant (Fig. S2). Although this assay allowed us to monitor mutations in real-time in the biofilm, this assay is also biased towards mutations that restore the frame of translation. To know the true impact of *dnaE2* on bacterial mutagenesis, the sequencing of the genomic DNA from biofilm culture would be the most appropriate assay but, was not performed in the present study.

To understand the significance of *dnaE2* expression in biofilm cultures, we competed the mutant strain with the parental strain by co-culturing in biofilm condition. While the mutant did not display significant growth defect when cultured individually in planktonic state, there was a drastic reduction in the population density of mutant in comparison to parental strain after 5th passage (Fig. 4A panels ii-iii and 4C). Since the contribution of DnaE2 to mutagenesis was modest we reasoned that additional roles of DnaE2 could be important for bacterial survival in biofilm. Previous studies have indicated that in *E. coli* expression of Pol IV impacted the DNA synthesis^39^. Oxidative DNA damage results in the formation of 8-oxo-G hindering the movement of replisome and may require the participation of translesion synthesis activity of DnaE2. However, the association of DnaE2 on the genomic DNA during translesion synthesis may create a “replication conflict” with the house keeping DnaE1 polymerase resulting in reduced replication rates and growth rate. Indeed, when the bacterial growth in planktonic and biofilm phases were compared, we observed attenuated bacterial growth in biofilm when DnaE2 was expressed (Fig. 2B and Fig. 5A-B). To further validate this hypothesis we employed a different strategy and treated WT and Δ*dnaE2* strains to Cip (an antibiotic that induces strong expression of *dnaE2*) and observed that the growth of WT strain was significantly slower than a similarly treated Δ*dnaE2* (Fig. 5C and 5D). Viability with reduced multiplication is the hallmark of persister phenotype in bacteria ^56,57^ and the replication conflict emanating from the expression of *dnaE2* could be one of the multiple factors triggering mycobacterial persistence in biofilm.

Till recently, the significance of mycobacterial biofilm was under appreciated as it was mainly studied in the *in vitro* context. However, occurrence of *M. tuberculosis* biofilms *in vivo* adds a new perspective on the ability of the pathogen’s ability to survive within the host ^19^. Antibiotic therapy and the evolution of clinically relevant drug resistance in *M. tuberculosis* is often studied together as a “cause and effect” paradigm. Our study highlights that the biofilms serve as a fertile niche for mycobacterial persistence and accelerating evolution in an antibiotic independent fashion directed by the expression of mutasome.

## Materials and Methods

### Bacterial strains and culture conditions

The list of strains, plasmids and oligonucleotides used in the study are presented in Supplementary Tables 1-3. Glycerol stocks of strains of *M. smegmatis* were revived on 7H10 Middlebrook supplemented with 0.5% v/v glycerol and 0.05% v/v Tween 80 plates, containing appropriate antibiotics. For culture in liquid condition, Middlebrook 7H9 medium supplemented with 0.1% v/v glycerol and 0.05%, Tween 80 (7H9T), along with appropriate antibiotics was used. For biofilm culturing, Middlebrook 7H9 medium supplemented with 0.5% v/v glycerol was used. When required, *M. smegmatis* cultures were selected by incorporating kanamycin (K) (50 µg/mL), apramycin (A) (25 µg/mL), and hygromycin (H) (50 µg/mL). For cloning and plasmid DNA manipulation, *E. coli* TG1 strains were used. *E. coli* strains were cultured in Luria Bertani (LB) broth or LB agar at 37 °C. For selection in *E. coli*, antibiotics with following concentrations kanamycin (50 µg/mL), apramycin (50 µg/mL), hygromycin (150 µg/mL) and ampicillin (100 µg/mL) were used. All media components were procured from Difco (Maryland, USA). Unless mentioned, all chemicals and reagents were procured from Sigma Aldrich (Missouri, USA).

### Expression analysis of reactive oxygen species (ROS) in mycobacterial biofilm

Freshly revived *M. smegmatis* mc^2^ 155 wild type (WT) on 7H10 plate was seeded for the biofilm in 7H9 biofilm media. A bacterial suspension corresponding to OD_600_ of 2.0 was prepared in biofilm medium. The bacterial suspension was diluted 1:100 in triplicates into individual cell of a 24 well sterile plate (Nunc, Denmark) containing 2 mL of biofilm medium. The biofilm culture was incubated at 30 °C in a static incubator. Along with this, a planktonic culture was set up in 7H9T media with initial OD_600_ corresponding to 0.05-0.1. This was cultured at 37 °C in a shaker incubator at 175 rpm. Planktonic culture in its early log phase (OD_600_ ∼0.3-0.4) was used as the negative control. After 2^nd^ and 4^th^ days of incubation, biofilm was disrupted by vigorous pipetting. The cells stuck to the wells were collected by washing the wells with 7H9 containing 0.2% Tween 80 (7H9-high Tween media). The bacterial cells were harvested by centrifuging at 13000 rpm for 1 minute. The harvested bacterial pellets were washed twice with 7H9-high Tween 80 media and resuspended in 90 µL 1X PBS with 0.2% Tween 80 (PBS-high Tween). The dried powder of ROS indicator dye CM-H_2_DCFDA (Invitrogen, California USA) was dissolved in 100 µL 100% DMSO to obtain a ∼80 mM stock solution. This solution was added to the bacterial suspension in PBST to a final concentration of 8 µM. The bacterial samples were incubated in the dark for 30 min at 37 °C. The samples and the planktonic culture controls were concentrated by mild centrifugation, after which excess broth/solution was discarded. The bacterial suspension with the residual media/broth was mounted on glass slides with ProLong Glass Antifade Mountant (Thermo-Fisher Scientific, Oregon USA) and subjected to Confocal Laser Scanning Microscopic analysis (CLSM). The images were acquired by Olympus FLUOVIEW FV3000 equipped with a 60X oil objective lens, through CCD camera connected to the FV3000 in-built software. Fields were selected randomly, and images were taken using Z-stacking with 60X oil immersion at the resolution of 512×512 pixels. The lens has a numerical aperture of 1.42. The images for the oxidized state (stained) were captured by exciting with 488 nm laser collecting the emission using 525/50 nm filter. The post-processing and analysis of the images were done using Olympus cellSens software platform. In the cellSens software, the images which were acquired with 96 ppi were projected in maximized view. For better image representation, the Brightness/Contrast of green was adjusted to 1556 from 4095. The images were saved as “.tiff” files.

### Estimation of NADH/NAD+ in biofilm cells

Freshly revived *M. smegmatis* mc^2^155 strain transformed with pMV762-Peredox-mCherry strain was seeded for biofilm as described above onto 7H9T medium containing hygromycin (7H9H). To image the biofilm, a sterile round glass coverslip sterile (12 mm, Blue Star, India) was dispensed to the well of 24 well sterile plate. After 2^nd^ and 4^th^ day, the media beneath the biofilm was aspirated out slowly and the biofilm was allowed to settle on the coverslip. Early log-phase of planktonic culture of these same strains (OD_600_ of ∼0.3-0.4) was kept as the control. The biofilm containing coverslips were mounted on glass slides with ProLong Glass Antifade Mountant. Along with it 100-200 µL of the control samples were centrifuged to harvest the cells and it was mounted on glass slides with the ProLong glass Mountant. Images were acquired using with Nikon *Eclipse Ti* microscope equipped with 60X oil immersion objective and Z-stacking feature, at the resolution of 512×512 pixels with the numerical aperture of 1.49, through CCD camera connected to the Nikon A16 computer system running NIS elements software. Images were acquired using the Nikon *EclipseTi* microscope equipped with 60X oil immersion objective, Z-stacking with resolution of 512×512 pixels. The images were captured by exciting with 405 nm laser and collecting the emission using 525/50 nm filter for T-sapphire, and exciting at 561 nm laser and emission at 625/50 nm filter for mCherry. The images were assigned a pseudo color for presentation of the NADH:NAD^+^ levels in individual cells. The post-processing and analysis were done by Nikon NIS elements version 4.00.04 software, and images were saved as “.tiff” format.

### Analysis of temporal expression of *recA* in mycobacterial biofilm

*M. smegmatis* mc^2^155 strains harboring pMV262 vector with *P*_*recA*_*∼*mClover was revived freshly on 7H10K plates. Biofilm was seeded in the 7H9 biofilm media with glass cover slip as described earlier. After 2^nd^, 4^th^ and 8^th^ day of incubation, the biofilm containing coverslips were mounted on glass slides with ProLong Glass Antifade Mountant. Images were acquired using Nikon *Eclipse Ti* microscope with appropriate laser for mClover (λ_ex_405nm / λ_em_525/50 nm) as explained above. The post-processing and analysis were done by Nikon NIS elements version

4.00.04 software. For better image representation, the Brightness/Contrast of green was adjusted to 1000 from 4095. Images were converted to “.tiff” format.

### Expression profiling of *recA*-dependent *dnaE2* expression in biofilm

*M. smegmatis* mc^2^155 harboring dual reporter, i.e., pMV262 vector with *P*_*recA*_*∼*mClover and *P*_*dnaE2*_*∼*mCherry together, was revived on a fresh 7H10K plate Biofilm was seeded in the 7H9 biofilm media with glass cover slip as described earlier. After 2^nd^, 4^th^ and 8^th^ day of incubation, the biofilm containing coverslips were mounted on glass slides with ProLong Glass Antifade Mountant. Images of the biofilm were acquired using Nikon *Eclipse Ti* microscope equipped with 60X oil immersion objective. Appropriate lasers for mClover (λ_ex_405nm / λ_em_525/50 nm) and mCherry (λ_ex_561nm / λ_em_625/50 nm) were used. The post-processing and analysis were done by Nikon NIS elements version 4.00.04 software. For better image representation, the Brightness/Contrast of both red and green were adjusted to 500 from 4095. Images were converted to “.tiff” format.

### Expression analysis of *dnaE2* accessory protein *imuA* in biofilm

*M. smegmatis* mc^2^ 155 harboring pMV262 vector with *P*_*imuA*_*∼*mCherry was subjected to biofilm formation as described previously and the expression of *imuA* was observed after 2^nd^, 4^th^ and 6^th^ day of incubation. Images with mCherry (λ_ex_561nm / λ_em_625/50 nm) were acquired by Olympus FLUOVIEW FV3000 equipped with a 60X oil immersion objective lens. The images post-processing and analysis of the images were done using Olympus cellSens software platform. In the cellSens software, the images which were acquired with 96 ppi were projected in maximized view. For better image representation, the Brightness/Contrast for red was adjusted to 1250 from 4095 and gamma value to 0.8 from 1. The images were saved as “.tiff” files.

### Time-course analysis of *dnaE2* expression during the biofilm recovery phase

*M. smegmatis* harboring the pMV262*P*_*hsp*_*∼*mClover and *P*_*dnaE2*_*∼*mCherry dual reporter strain was subjected to biofilm formation as described earlier without glass coverslip. After 6 days, the culture was harvested at 13,000 rpm for 1 min and washed twice with 7H9 high Tween 80 (2% Tween 80) media. The washed bacterial pellet was resuspended in fresh 7H9K and incubated at 37 °C at 175 rpm. An aliquot of this harvested culture was concentrated and mounted on a glass slide immediately with ProLong Glass Antifade Mountant and considered as 0 h after recovery. Subsequently, a sample from the recovery was analyzed after 3 and 6 h for the expression of dual reporters by CLSM. The images were acquired by Olympus FLUOVIEW FV3000 using appropriate lasers for mClover (λ_ex_405nm / λ_em_525/50 nm) and mCherry (λ_ex_561nm / λ_em_625/50 nm). Fields were selected randomly, and were captured using Z-stacking with 60X oil immersion at the resolution of 512×512 pixels. The post-processing and analysis of the images were done using Olympus cellSens software platform. In the cellSens software, the images which were acquired with 96 ppi were projected in maximized view and the images were saved as “.tiff” files.

### Determination of viability of bacterial cells in biofilm culture by live/dead staining

Freshly revived *M. smegmatis* mc^2^155 was subjected to biofilm formation and planktonic cultures as described previously. After 6 day of incubation the biofilm was disrupted and washed twice with 7H9 high Tween 80 (0.2% vol/vol) media and harvested the cell. For planktonic culture, cells were harvested from the mid-log phase culture. For Live/Dead staining, the bacterial pellets were resuspended in 1 mL of 150 mM NaCl solution containing 0.2% Tween 80 and stained with Live/Dead staining solution for 15 min at room temperature in the dark. The staining solution was prepared by mixing Syto9 and propidium iodide in 1:1 ratio which were reconstituted in DMSO (Invitrogen, California, USA). The samples were concentrated by mild centrifugation and mounted on glass slides with ProLong Glass Antifade Mountant, as described earlier. The images were acquired by Olympus FLUOVIEW FV3000 confocal microscope using Z-stacking with 60X oil immersion. The Syto9 (live) images were captured with λ_ex_ 405nm/λ_em_ 525/50 nm. PI (dead) images were captured with λ_ex_ 561nm / λ_em_ 625/50 nm. The post-processing and analysis of the images were done using Olympus cellSens software platform, as mentioned above. For better image representation, the Brightness/Contrast of green was adjusted to 1556 from 4095. The images were saved in “.tiff” format.

### Determination of mutation rate in biofilm and planktonic cultures of WT, *ΔdnaE2* and compliment strains of *M. smegmatis*

*M. smegmatis* mc^2^155 with empty vector (WT), *ΔdnaE2*, and *ΔdnaE2::pdnaE2* strains were freshly revived on 7H10A. The cultures made into bacterial suspension of OD_600_ ∼2.0 and 20 µL was added to 2 mL 7H9 biofilm media in a sterile flat bottom 24-well plate for biofilm and in cultures tubes containing 7H9T media for planktonic with the starting density of OD_600_∼0.02 for both culturing systems. For each strain ten replicates were kept in both planktonic and biofilm condition. The biofilm culture was incubated at 30 °C in a static incubator and planktonic culture at 37 °C in a shaker incubator at 175 rpm. Once the planktonic culture reached around ∼0.4-0.6 OD_600_, an aliquot (50 μL) was taken out from each replicate, proceeded with the tenfold serial dilution till 10^−4^, to enumerate the viable counts. The remaining culture was harvested in a sterile 2 mL microcentrifuge tube at 13,000 rpm for 1 min at room temperature. The supernatant was discarded, and the pellet was resuspended in 0.1 mL of 7H9 medium. This suspension was spread-plated on 7H10 plates containing 100 µg/mL rifampicin (Rif) or 2.5 µg/mL ciprofloxacin (Cip) to enumerate the mutants in the culture. After 6^th^ day of incubation biofilm was disrupted and cells were harvested. The harvested cells were washed twice with 7H9 high Tween 80 media. Then the same procedure as of planktonic culture was followed to enumerate viable counts and mutant counts in the biofilm culture. The plates were incubated at 37 °C, and viable count were recorded after 3 days, and Rif^R^ mutants and Cip^R^ mutants were recorded after 5 days and 7 days respectively. The mutation frequency was determined by calculating the number of mutants obtained on Rif or Cip plate with the corresponding viable count and statistical analysis and the graph plotting was done using GraphPad Prism^®^software version 8.

### Competition assay in biofilm

*M. smegmatis* mc^2^155 with *P*_*hsp*_∼mClover (*WT*) and Δ*dnaE2 with P*_*hsp*_∼mCherry reporter strains were freshly revived on 7H10K. For each strain a bacterial suspension of OD_600_∼ 2 was prepared. The competition experiment was initiated by mixing competing strains i.e., the WT with *P*_*hsp*_∼mClover v/s Δ*dnaE2* with *P*_*hsp*_∼mCherry in 1:1 ratio with OD_600_∼ 0.02 each. The final density of the biofilm corresponding to OD_600_∼ 0.02 in 2 mL 7H9K biofilm media in a sterile 24-well flat bottom plate. This 1:1 mix was considered as “0^th^ passage”. The cultures were incubated for 6 days at 30 °C static condition to get the “1^st^ passage”. From this 100 μL-200 μL of homogenous bacterial suspension was plated on 7H10K agar. From the plate, biofilms were seeded with bacterial inoculum corresponds to OD_600_∼ 0.02. The seeding of biofilm process followed by the subsequent sub culturing process was repeated until the 6th passage is obtained. A quadrant streaking was done at 0^th^ and 6^th^ passage on 7H10K plate to get isolated colonies for patching on 7H10HK media as well as in 7H10K media, to enumerate the number of Δ*dnaE2 and* WT colonies respectively. At the end of every passage, 100-200 µl aliquots of the culture were taken, and processed for the microscopic analysis. Images were acquired using Nikon *Eclipse Ti* with appropriate lasers for mClover (λ_ex_405nm / λ_em_525/50 nm) and mCherry (λ_ex_561nm / λ_em_625/50 nm). The post-processing and analysis were done by Nikon NIS elements version 4.00.04 software. The area of focus was zoomed in by 3X for better representation. Images were converted to “.tiff” format.

### Growth kinetics of *M. smegmatis* in planktonic and biofilm system

*M. smegmatis* mc^2^155 WT was freshly revived on 7H10. A bacterial suspension with density corresponding to the OD_600_ ∼ 2.0 was prepared. The seeding of both planktonic and biofilm cultures was done as described earlier with the starting OD_600_ as 0.02. The biofilm culture was incubated at 30 °C in a static incubator and planktonic culture at 37°C at 175 rpm. The observation points were 0 h, 2, 4 and 6 days with three replicates kept for each time point for both planktonic and biofilm samples. At each time point biofilm was disrupted, the cells were harvested and washed with 7H9 high Tween 80 media and the samples were vortexed to get uniform suspension and measured at OD_600_ in a Spectrophotometer (Thermo Fisher scientific, USA). Along with the optical density, the total viable counts were enumerated by serial dilution plating method. The obtained bacterial growth profile through spectrophotometer and the viable counts of both the condition were plotted into a graphical representation using GraphPad Prism^®^software version 8.

### RNA sequencing of the biofilm and the planktonic system

*M. smegmatis* mc^2^155 WT strain was freshly revived on 7H10 plate. The biofilm and planktonic cultures were seeded in the 7H9 biofilm media and 7H9 media respectively, as described earlier. For biofilm recovery samples, the biofilm was disrupted and wash twice with high tween 7H9 and incubated in 7H9 for 3 h. The cells were harvested after the respective incubation time. The harvested pellets from each condition were immediately frozen using the liquid nitrogen and stored at -80°C. The pellets were resuspended in 1 mL TRI reagent with gentle pipetting on ice. This suspension was transferred into a sterile 2 mL bead beating tube containing 0.5mm Zirconia-Silicate beads (Biospec, USA). The bead beating was carried on in a Mini-beadbeater (BioSpec, USA). The bead beating was done in 2 cycles of 1 minute each with an intermittent cooling on ice for 2 min between the cycles. These tubes were centrifuged at 13000 rpm for 10 min at 4 °C in an Eppendorf 5804 R centrifuge with microcentrifuge tube rotor. The supernatant from the cell lysate was transferred to a new sterile tube leaving behind cell debris and beads. Another centrifugation was done at 14000 rpm for 10 min at 4 °C and the supernatant was transferred to a sterile microcentrifuge tube. To this tube, chloroform was added in 1/5^th^ volume of the supernatant. The aqueous layer and the chloroform was gently mixed by rotating the tube and incubated for 10 min at room temperature. This mix was centrifuged at 14000 rpm for 10 min at 4 °C. The aqueous layer was carefully transferred to a new 1.7 mL microcentrifuge tube the RNA was precipitated by adding 600 µL of isopropanol with gentle mixing. This was kept at -20 °C for an overnight incubation. After the incubation the vial with precipitation was centrifuged at 13000 rpm at 4 °C for 20 min and remove the supernatant with carefully without disturbing the minute pellet. This pellet was washed with 80% ethanol prepared with DEPC treated water, centrifuged at 13000 rpm at 4 °C. The ethanol was removed. The centrifugation was repeated one more time to remove excess ethanol. The pellet was then air dried on a clean surface for 2-3 min. The pellet was resuspended in 88 µL of DEPC treated water and made to dissolve on ice. The dissolved RNA was added with 10 µL of DNase buffer and 2 µL of DNase enzyme (ThermoFisher scientific, Massachusetts, United States), and incubated at 37 °C for 30 min. The RNA was further purified using RNA isolation kit by Machery-Nagel (Germany) according to manufacturer’s protocol. The quality of RNA was checked by both UV spectrometry and agarose gel electrophoresis.

### Library preparation

Briefly, 1µg of total RNA from each sample (three groups, with duplicate sample in each group) was depleted of bacterial rRNA using NEBNext® rRNA depletion kit (Cat. no. E7850X, New England Biolabs, Ipswich, MA,USA), according to the manufacturer’s protocol. This included a DNaseI digestion step. The rRNA depleted RNA was purified using RNA purification beads as instructed. The NEBNext® Ultra II Directional RNA Library prep kit for Illumina (Cat. no. E7760L), was used to construct double-stranded cDNA libraries from the rRNA depleted RNA, using RNA fragmentation, first strand cDNA synthesis with random primers, second strand cDNA synthesis, end repair of double stranded cDNA, adapter ligation, removal of excess adapter using sample purification beads, PCR enrichment of adapter ligated DNA and clean-up of PCR products as instructed by the manufacturer. The cleaned libraries were quantitated on Qubit flurometer (Thermo Fisher Scientific, Waltham, MA,USA) and appropriate dilutions loaded on a Tapestation 4200 High sensitivity DS1000 screen tape (Agilent Technologies, Santa Clara, CA,USA) to determine the size range of the fragments and the average library size.

### Sequencing and data processing

Libraries were diluted to 4 nM, pooled, spiked with 5% PhiX pre-made library from Illumina and loaded on a Miseq v3 kit (Illumina Inc., San Diego,CA,USA). Sequencing was performed for 1×150 cycles. The original raw data from Illumina Miseq were transformed to sequenced reads by base calling. The Raw data were recorded in FASTQ files and the reads per sample ranged from 3497155 to 5051470. The quality of the reads was assessed using *FastQC v 0*.*11*.*3* ^60^ before proceeding for the downstream analysis. The reads were trimmed using *Fastpv 0*.*20*.*1* ^58^ when the quality of the bases dropped below 30, and adapters were removed. Reads were rechecked for improvement in quality before proceeding further. A transcriptome index was prepared using the cDNA of reference assembly *M. smegmatis* mc^2^155 obtained from NCBI (https://www.ncbi.nlm.nih.gov/assembly/GCF_000015005.1/). The quality passed reads were mapped onto the reference transcriptome and were quantified with *Kallisto v0*.*46*.*2* ^59^. The aligned reads ranged from 85.39% -92.35%.

### Bioinformatic analysis

The *Degust v4*.*1*.*1 web-tool* ^61^ was used for differential expression analysis. Only genes with count per million (CPM) ≥1 were analysed further. Genes were filtered based on false discovery rate cut-off (FDR) ≤0.05 and minimum expression fold change (FC) ≥2. The sequence reads has been deposited in the SRA server under the project name **PRJNA888550**.

## Acknowledgement

The authors thank Dr. R Ajay Kumar, Rajiv Gandhi Centre for Biotechnology, for sharing resources and lab space. The authors thank Prof. Umesh Varshney and lab members Dr. Shivjee and Shashank Aroli, Indian Institute of Science, for sharing the *recA*::*kan* strain and allowing the use of Bioscreen C instrument for growth curve analysis. Dr. Ashwani Kumar, Institute of Microbial Technology and Dr. Markus A Seegur, University of Zurich for sharing strains and plasmids used in the study. The authors thank Dr. Saravanan Matheshwaran, Indian Institute of Technology, Kanpur, Dr. Vijay Srinivasan, Microverse Cluster Jena, Dr Garima Khare, University of Delhi and Dr. Sabari, Indian Institute for Science Education and Research, Thiruvananthapuram for critical reading and valuable suggestions during the preparation of the manuscript. The Central Imaging facility, Rajiv Gandhi Centre for Biotechnology and miBiomeTheraputics LLB (Mumbai) are acknowledged for the assistance in microscopy and RNA sequencing analysis respectively. The schematics were generated using Library of Science and Medical Illustrations available at www.somersault1824.com. KK acknowledges the financial support from Ramalingaswami Re-entry fellowship from Department of Biotechnology, Government of India, intramural funds from Rajiv Gandhi Centre for Biotechnology and the extramural grant from Science and Engineering Board (CRG/2018/001209), Government of India.

## References

1. Hall-Stoodley, L., Costerton, J. W. & Stoodley, P. Bacterial biofilms: from the Natural environment to infectious diseases. Nat Rev Microbiol 2, 95–108 (2004).

2. Hansen, S. K., Rainey, P. B., Haagensen, J. A. J. & Molin, S. Evolution of species interactions in a biofilm community. Nature 445, 533–536 (2007).

3. Steenackers, H. P., Parijs, I., Dubey, A., Foster, K. R. & Vanderleyden, J. Experimental evolution in biofilm populations. Fems Microbiol Rev 40, 373–397 (2016).

4. Pesavento, C. & Hengge, R. Bacterial nucleotide-based second messengers. Curr Opin Microbiol 12, 170–176 (2009).

5. Camilli, A. & Bassler, B. L. Bacterial Small-Molecule Signaling Pathways. Science 311, 1113–1116 (2006).

6. Branda, S. S., Vik, Å., Friedman, L. & Kolter, R. Biofilms: the matrix revisited. Trends Microbiol 13, 20–26 (2005).

7. López, D., Vlamakis, H. & Kolter, R. Biofilms. Cold Spring Harbor Perspectives in Biology 2, a000398 (2010).

8. Vlamakis, H., Chai, Y., Beauregard, P., Losick, R. & Kolter, R. Sticking together: building a biofilm the Bacillus subtilis way. Nature Reviews Microbiology 11, 157–168 (2013).

9. Kolter, R., Vlamakis, H. & Gestel, J. van. Division of Labor in Biofilms: the Ecology of Cell Differentiation. Microbiology Spectrum 3, 1–24 (2015).

10. García-Betancur, J. C. & Lopez, D. Cell Heterogeneity in Staphylococcal Communities. J Mol Biol 431, 4699–4711 (2019).

11. Williamson, K. S. et al. Heterogeneity in Pseudomonas aeruginosa biofilms includes expression of ribosome hibernation factors in the antibiotic-tolerant subpopulation and hypoxia-induced stress response in the metabolically active population. J Bacteriol 194, 2062–73 (2012).

12. Wimpenny, J., Manz, W. & Szewzyk, U. Heterogeneity in biofilms. FEMS Microbiol Rev 24, 661–671 (2000).

13. Stewart, P. S. & Franklin, M. J. Physiological heterogeneity in biofilms. Nature Reviews Microbiology 6, 199–210 (2008).

14. Sadiq, F. A. et al. Phenotypic and genetic heterogeneity within biofilms with particular emphasis on persistence and antimicrobial tolerance. Future Microbiol 12, 1087–1107 (2017).

15. Donlan, R. M. Biofilm Formation: A Clinically Relevant Microbiological Process. Clin Infect Dis 33, 1387–1392 (2001).

16. Schulze, A., Mitterer, F., Pombo, J. P. & Schild, S. Biofilms by bacterial human pathogens: Clinical relevance – development, composition and regulation – therapeutical strategies. Microb Cell 8, 28 (2021).

17. Ojha, A. et al. GroEL1: A Dedicated Chaperone Involved in Mycolic Acid Biosynthesis during Biofilm Formation in Mycobacteria. Cell 123, 861–873 (2005).

18. Trivedi, A., Mavi, P. S., Bhatt, D. & Kumar, A. Thiol reductive stress induces cellulose-anchored biofilm formation in Mycobacterium tuberculosis. Nat Commun 7, 11392 (2016).

19. Chakraborty, P., Bajeli, S., Kaushal, D., Radotra, B. D. & Kumar, A. Biofilm formation in the lung contributes to virulence and drug tolerance of Mycobacterium tuberculosis. Nat Commun 12, 1606 (2021).

20. Recht, J. & Kolter, R. Glycopeptidolipid Acetylation Affects Sliding Motility and Biofilm Formation in Mycobacterium smegmatis. J Bacteriol 183, 5718–5724 (2001).

21. Recht, J., Martínez, A., Torello, S. & Kolter, R. Genetic Analysis of Sliding Motility inMycobacterium smegmatis. J Bacteriol 182, 4348–4351 (2000).

22. Yang, Y. et al. Defining a temporal order of genetic requirements for development of mycobacterial biofilms. Mol Microbiol 105, 794–809 (2017).

23. Ojha, A. & Hatfull, G. F. The role of iron in Mycobacterium smegmatis biofilm formation: the exochelin siderophore is essential in limiting iron conditions for biofilm formation but not for planktonic growth. Mol Microbiol 66, 468–483 (2007).

24. Yang, Y., Richards, J. P., Gundrum, J. & Ojha, A. K. GlnR Activation Induces Peroxide Resistance in Mycobacterial Biofilms. Front Microbiol 9, 1428 (2018).

25. Ojha, A. K. et al. Growth of Mycobacterium tuberculosis biofilms containing free mycolic acids and harbouring drug-tolerant bacteria. Mol Microbiol 69, 164–74 (2008).

26. Rose, S. J., Babrak, L. M. & Bermudez, L. E. Mycobacterium avium Possesses Extracellular DNA that Contributes to Biofilm Formation, Structural Integrity, and Tolerance to Antibiotics. Plos One 10, e0128772 (2015).

27. Penterman, J. et al. Rapid Evolution of Culture-Impaired Bacteria during Adaptation to Biofilm Growth. CellReports 6, 293–300 (2014).

28. Driffield, K., Miller, K., Bostock, J. M., O’Neill, A. J. & Chopra, I. Increased mutability of Pseudomonas aeruginosa in biofilms. J Antimicrob Chemother 61, 1053–6 (2008).

29. Ryder, V. J., Chopra, I. & O’Neill, A. J. Increased Mutability of Staphylococci in Biofilms as a Consequence of Oxidative Stress. Plos One 7, e47695 (2012).

30. Conibear, T. C. R., Collins, S. L. & Webb, J. S. Role of mutation in Pseudomonas aeruginosa biofilm development. Plos One 4, e6289 (2009).

31. Boshoff, H. I. M., Reed, M. B., Barry, C. E. & Mizrahi, V. DnaE2 polymerase contributes to in vivo survival and the emergence of drug resistance in Mycobacterium tuberculosis. Cell 113, 183–193 (2003).

32. Warner, D. F. et al. Essential roles for imuA’- and imuB-encoded accessory factors in DnaE2-dependent mutagenesis in Mycobacterium tuberculosis. Proceedings of the National Academy of Sciences of the United States of America 107, 13093–13098 (2010).

33. Bhat, S. A., Iqbal, I. K. & Kumar, A. Imaging the NADH:NAD+ Homeostasis for Understanding the Metabolic Response of Mycobacterium to Physiologically Relevant Stresses. Front Cell Infect Mi 6, 145 (2016).

34. Murphy, M. P. How mitochondria produce reactive oxygen species. Biochem J 417, 1–13 (2009).

35. Xie, N. et al. NAD+ metabolism: pathophysiologic mechanisms and therapeutic potential. Signal Transduct Target Ther 5, 227 (2020).

36. Imlay, J. A. The molecular mechanisms and physiological consequences of oxidative stress: lessons from a model bacterium. Nat Rev Microbiol 11, 443–454 (2013).

37. Zhou, W. et al. A Feedback Regulatory Loop Containing McdR and WhiB2 Controls Cell Division and DNA Repair in Mycobacteria. Mbio 13, e03343–21 (2022).

38. Salini, S. et al. The Error-Prone Polymerase DnaE2 Mediates the Evolution of Antibiotic Resistance in Persister Mycobacterial Cells. Antimicrob Agents Ch 66, e01773–21 (2022).

39. Indiani, C., Langston, L. D., Yurieva, O., Goodman, M. F. & O’Donnell, M. Translesion DNA polymerases remodel the replisome and alter the speed of the replicative helicase. P Natl Acad Sci Usa 106, 6031–8 (2009).

40. Oliver, J. D. The viable but nonculturable state in bacteria. J Microbiol Seoul Korea 43 Spec No, 93–100 (2005).

41. Ayrapetyan, M., Williams, T. & Oliver, J. D. Relationship between the Viable but Nonculturable State and Antibiotic Persister Cells. J Bacteriol 200, 549–15 (2018).

42. Gotoh, H., Kasaraneni, N., Devineni, N., Dallo, S. F. & Weitao, T. SOS involvement in stress-inducible biofilm formation. Biofouling 26, 603–611 (2010).

43. Oh, E., Kim, J.-C. & Jeon, B. Stimulation of biofilm formation by oxidative stress in Campylobacter jejuni under aerobic conditions. Virulence 7, 1–6 (2016).

44. Cepas, V. et al. Relationship Between Biofilm Formation and Antimicrobial Resistance in Gram-Negative Bacteria. Microb Drug Resist 25, 72–79 (2019).

45. Cáp, M., Váchová, L. & Palková, Z. Reactive Oxygen Species in the Signaling and Adaptation of Multicellular Microbial Communities. Oxid Med Cell Longev 2012, 976753 (2012).

46. Gambino, M. & Cappitelli, F. Mini-review: Biofilm responses to oxidative stress. Biofouling 32, 167–178 (2016).

47. Geier, H., Mostowy, S., Cangelosi, G. A., Behr, M. A. & Ford, T. E. Autoinducer-2 Triggers the Oxidative Stress Response in Mycobacterium avium, Leading to Biofilm Formation. Appl Environ Microb 74, 1798–1804 (2008).

48. Kreth, J., Vu, H., Zhang, Y. & Herzberg, M. C. Characterization of Hydrogen Peroxide-Induced DNA Release by Streptococcus sanguinis and Streptococcus gordonii. J Bacteriol 191, 6281–6291 (2009).

49. Aldecoa, A. L. I. de, Zafra, O. & González-Pastor, J. E. Mechanisms and Regulation of Extracellular DNA Release and Its Biological Roles in Microbial Communities. Front Microbiol 8, 1390 (2017).

50. Boles, B. R. & Singh, P. K. Endogenous oxidative stress produces diversity and adaptability in biofilm communities. Proc National Acad Sci 105, 12503–12508 (2008).

51. Bernier, S. P. et al. Starvation, Together with the SOS Response, Mediates High Biofilm-Specific Tolerance to the Fluoroquinolone Ofloxacin. Plos Genet 9, e1003144–14 (2013).

52. García-Castillo, M. et al. Stationary biofilm growth normalizes mutation frequencies and mutant prevention concentrations in Pseudomonas aeruginosa from cystic fibrosis patients. Clin Microbiol Infec 17, 704–711 (2011).

53. Brandis, G., Pietsch, F., Alemayehu, R. & Hughes, D. Comprehensive phenotypic characterization of rifampicin resistance mutations in Salmonella provides insight into the evolution of resistance in Mycobacterium tuberculosis. J Antimicrob Chemoth 70, 680–685 (2015).

54. Melnyk, A. H., Wong, A. & Kassen, R. The fitness costs of antibiotic resistance mutations. Evol Appl 8, 273–283 (2015).

55. Mokrzan, E. M. et al. Nontypeable Haemophilus influenzae newly released (NRel) from biofilms by antibody-mediated dispersal versus antibody-mediated disruption are phenotypically distinct. Biofilm 2, 100039 (2020).

56. Balaban, N. Q., Merrin, J., Chait, R., Kowalik, L. & Leibler, S. Bacterial Persistence as a Phenotypic Switch. Science 305, 1622–1625 (2004).

57. Radzikowski, J. L., Schramke, H. & Heinemann, M. Bacterial persistence from a system-level perspective. Curr Opin Biotech 46, 98–105 (2017).

58. Chen, S., Zhou, Y., Chen, Y. & Gu, J. fastp: an ultra-fast all-in-one FASTQ preprocessor. Bioinformatics 34, i884–i890 (2018).

59. Bray, N. L., Pimentel, H., Melsted, P. & Pachter, L. Near-optimal probabilistic RNA-seq quantification. Nat Biotechnol 34, 525–527 (2016).

60. Andrews, S. (2010). FastQC: A Quality Control Tool for High Throughput Sequence Data [Online]. Available online at: http://www.bioinformatics.babraham.ac.uk/projects/fastqc/

61. David R. Powell. Degust: interactive RNA-seq analysis, DOI: 10.5281/zenodo.3258932

