## Supplementary data and table for "*dnaE2* expression promotes genetic diversity and bacterial persistence in mycobacterial biofilms"

Running title: Mutagenesis in mycobacterial biofilm

Keywords: Mycobacteria, Biofilms, Mutagenesis, Persistence, Fitness defect, Bacterial evolution

**Supplementary methods**

**Determination of *recA* dependent *dnaE2* expression with *∆recA* KO**

*M smegmatis ∆recA::kan* strain with *P_dnaE2_*~mCherry-hyg plasmid was freshly revived on 7H10HK plate. Biofilm was seeded in the 7H9 biofilm media with glass cover slip as described earlier. After 2^nd^, 4^th^ and 8^th^ day of incubation, the biofilm containing coverslips were mounted on glass slides with ProLong Glass Antifade Mountant. Images of the biofilm and the control samples were acquired using the appropriate filters for mCherry (λ_ex_561nm / λ_em_625/50 nm) in Nikon *Eclipse Ti* microscope, with 60X oil immersion objective. For better image representation, the Brightness/Contrast of both red and green were adjusted to 750 from 4095. Images were converted to ".tiff" format.

**Site directed mutagenesis of GFP to generate +1 frameshift**

Mycobacterial plasmid pFPV expressing stable version of GFP through the *P_hsp_* was used as template for generating non-fluorescent version of GFP. Oligonucleotides GFP+1Fp and GFP+1 Rp were designed to introduce an extra G at 313 position of GFP orf using the PCR based Q5 site directed mutagenesis (NEB) online tool. Briefly, 25 ng of pFPVGFP was incubated with 10 pmol each of GFP+1 Fp and GFP+1 Rp in a 25 µL containing 1X of Q5 Hot start high fidelity master mix. The PCR reaction was incubated at 98 ^o^C for 30 s followed by 24 cycles of incubation at 98 ^o^C for 15 s, 69 ^o^C for 30 s and 72 for 2 min 30 s. The PCR was processed according to manufacturer’s protocol and the final product DNA was transformed into *E*. *coli* TG1 competent cells and selected on LB agar containing kanamycin. Plasmid DNA from four isolated transformants was isolated and the presence of +1 frame shift mutation was confirmed by DNA sequencing and denoted as pFPVGFP*. Plasmid pFPVGFP* was introduced into *M. smegmatis* strains by electroporation for further analysis.

**GFP conversion Assay**

*M. smegmatis* mc^2^155 with *P_hsp_*~GFP* and Δ*dnaE2 with P_hsp_*~GFP* reporter strains were seeded for biofilm formation as mentioned earlier. After 6^th^ day of incubation, the biofilm was harvested and mounted on glass slide with ProLong Glass Antifade Mountant and subjected to CLSM. Images were acquired using with Nikon *Eclipse Ti* microscope equipped with 60X oil objective and Z-stacking feature with appropriate lasers for GFP (λ_ex_405nm / λ_em_525/50 nm), at the resolution of 512X512 pixels with the numerical aperture of 1.49, through CCD camera connected to the Nikon A16 computer system running NIS elements software. The post-processing and analysis were done by Nikon NIS elements version 4.00.04 software. And the images were converted to ".tiff" format.

**Supplementary Tables**

Table S1. List of bacterial strains

| **Strain** | **Relevant details** | **Reference** |
| --- | --- | --- |
| *E. coli* TG1 | Derivative strain of *Escherichia coli* JM101 |  |
| *M. smegmatis* mc^2^155 | High-efficiency transformation strain of *Mycobacterium smegmatis.* | 1 |
| *M. smegmatis*  *recA::kan* | *M. smegmatis* in which *recA* has been insertionally inactivated by insertion of kanamycin marker | 2 |
| mc^2^155-*P_recA_ reporter* | mc^2^155 strain transformed with *recA* reporter pMV262~mClover replicative plasmid and is resistant to kanamycin. | 3 |
| mc^2^155-Peredox | mc^2^155 strain transformed with the hygromycin resistant pMV762-Peredox-mCherry plasmid to monitor NADH/NAD^+^levels in bacterium. | 3 |
| mc^2^155-pMV262 ~*dual reporter* | mc^2^155 strain harbouring pMV262 expression vector with P*_recA_*~mClover and P*_dnaE2_*~mCherry together. | 3 |
| *recA::kan*  p*dnaE2*~mCherry *hyg* | mc^2^155 *recA* mutant strain harbouring the *dnaE2* reporter in hygromycin mycobacterial replicative plasmid. | 3 |
| Δ*dnaE2* | mc^2^155 *dnaE2*::*hyg^R^* strain transformed with p*att-FLAG* integrative plasmid and is resistant to both hygromycin and apramycin. | 3 |
| Δ*dnaE2*::*pdnaE2* | mc^2^155 *dnaE2*::*hyg^R^* strain transformed with p*att-dnaE2_UP* integrative plasmid and is resistant to both hygromycin and apramycin. | 3 |
| Δ*dnaE2~*pMVmCherry | mc^2^155 *dnaE2*::*hyg^R^* strain transformed with P*_hsp_*~mCherry replicative plasmid and is resistant to kanamycin and hygromycin. | This study |
| mc^2^155-pMV~mCherry | mc^2^155 strain transformed with pMV~mCherry replicative plasmid and is resistant to kanamycin | This study |
| mc^2^155-pMV~mClover | mc^2^155 strain transformed with pMV~mClover replicative plasmid and is resistant to kanamycin | This study |
| mc^2^155-*P_imuA_ reporter* | mc^2^155 strain transformed with *imuA* reporter pMV262~mCherry replicative plasmid and is resistant to kanamycin. | This study |
| mc^2^155-pMV262 ~*dual reporter* | mc^2^155 strain harbouring pMV262 expression vector with P*_recA_*~mClover and P*_dnaE2_*~mCherry | This study |
| mc^2^155-pFPV~GFP* | mc^2^ 155 strain transformed with kanamycin resistant pFPV~GFP* non fluorescent plasmid for GFP reversion assay | This study |
| Δ*dnaE2-* pFPV~GFP* | Δ*dnaE2* strain transformed with kanamycin resistant pFPV~GFP* non fluorescent plasmid for GFP reversion assay | This study |

**Table S2. Plasmid list and description**

| **Plasmid** | **Sequence/Relevant details** | **Reference** |
| --- | --- | --- |
| pJet1.2 | Blunt-end PCR cloning vector | Thermofisher Scientific |
| pMV~mCherry | Mycobacterial replicative plasmid with pAL5000 mycobacterial origin of replication encoding mCherry fluorescent protein from a constitutive *P_hsp_* promoter cloned between *BamH*I and *EcoR*I sites. The mCherry was designed to contain an in frame additional 12 amino acids corresponding to the *E. coli tmRNA* just before the stop-codon to destabilize the protein. The plasmid confers kanamycin resistance. | Kind gift from Dr. Marcela G. Rodriguez |
| p*dnaE2~*mCherry | Short half-life pMV~mCherry plasmid in which the *P_hsp_* promoter was replaced by digestion with *NotI* and *BamHI* restriction endonucleases with 300 bp region upstream to the *M. tuberculosis dnaE2* orf encompassing the *dnaE2* promoter (*P_dnaE2_*). | 3 |
| p*dnaE2*~mCherry *hyg* | Same as the above plasmid but the kanamycin marker is replaced with hygromycin resistance marker. | 3 |
| pMV*~*mClover | Mycobacterial replicative plasmid with pAL5000 mycobacterial *ori* expressing short half-life mClover fluorescent protein from a constitutive *P_hsp_* promoter cloned between *BamHI* and *EcoRI* sites. The mCherry was designed to contain an in frame additional 12 amino acids corresponding to the *E. coli tmRNA* just before the stop-codon. | 3 |
| pMV~dual reporter | pMV plasmid containing both the *P_dnaE2_~*mCherry and *P_hsp_~*mClover reporters in a single plasmid with kanamycin marker. | 3 |
| pMV~*recA*-dual reporter | pMV plasmid containing both the *P_dnaE2_~*mCherry and *P_recA_~*mClover reporters in a single plasmid with kanamycin marker. |  |
| p*recA~mClover* | Short half-life pMV~mClover plasmid in which the *P_hsp_* promoter was replaced by digestion with *NotI* and *BamHI* restriction endonucleases with 300 bp region upstream to the *M. smegmatis recA* orf encompassing the *recA* promoter (*P_recA_*). | 3 |
| *P_imuA_~*mCherry | Short half-life pMV~mCherry plasmid in which the P*_hsp_* promoter was replaced by digestion with *NotI* and *BamHI* restriction endonucleases with 300 bp region upstream to the *M. tuberculosis* *imuA* orf encompassing the *imuA* promoter (*P_imuA_*). | This study |
| pMV762-Peredox-mCherry | Mycobacterial plasmid expressing NADH:NAD^+^ ratio biosensor with hygromycin marker and pAL5000 origin of replication. | 3 |
| pFPV-GFP* | Mycobacterial plasmid pFPV-GFP was mutagenized to introduce a +1 frameshift in the GFP open reading frame at 313 position. The GFP is expressed through *P_hsp_* promoter and the plasmid contains pAL5000 origin of replication with kanamycin marker | This study |

**Table S3. List and sequence of oligonucleotides**

| **Name** | **Description** | **Sequence (5’-3’)** |
| --- | --- | --- |
| Msm*imuA*_300-Fp | *imuA* promoter 300 up of *M. smegmatis* for reporter. | GCGGCCGCcccatcccgaagcgggtt (NotI) |
| Msm*imuA*_300-Rp |  | GGATCCaccagcacctcccgcacc (BamHI) |
| GFP+1Fp | Introduces +1 frame shift in GFP at position 313 in the orf making it non-functional | CGTGCTGAAGTCAAGTTTGAAGGTGA |
| GFP+1Rp |  | TGTCTTGTAGTTCCCCGTCATCTT T |

**Reference**

1) Snapper S. B., Melton R. E., Mustafa S., Kieser T., Jacobs W. R. Jr. Isolation and characterization of efficient plasmid transformation mutants of *Mycobacterium smegmatis*. Mol Microbiol 1990; 4:1911–1919
2) Pau Biak Sang, Thiruneelakantan Srinath, Aravind Goud Patil, Eui-Jeon Woo, and Umesh Varshney. A unique uracil-DNA binding protein of the uracil DNA glycosylase superfamily. Nucleic Acids Res. 2015; 43(17): 8452–8463.
3) Salini S, Bhat SG, Naz S, Natesh R, Kumar RA, Nandicoori VK, Kurthkoti K. The Error-Prone Polymerase DnaE2 Mediates the Evolution of Antibiotic Resistance in Persister Mycobacterial Cells. Antimicrob Agents Chemother. 2022; 66(3):e0177321.
4) Shabir A. Bhat, Iram K. Iqbal, and Ashwani Kumar. Imaging the NADH:NAD+

Homeostasis for Understanding the Metabolic Response of Mycobacterium to Physiologically Relevant Stresses. Front Cell Infect Microbiology. 2016; 6, 145

**Legends to figures**

**Figure S1. RecA deletion reduces *dnaE2* expression in biofilm.** *M. smegmatis* *recA*::*kan* containing pMV262 *P_dnaE2_~*mCherry was seeded for biofilm with glass coverslips as mentioned before. At regular intervals the coverlip layered with biofilm was subjected to confocal microscopic analysis and fluorescent images were acquired. Representative image is presented.

**Figure S2. Deletion of *dnaE2* reduces mutation within biofilm. A)** DNA sequence chromatogram for the confirmation of generation of pFPVGFP* containing a+1 frame shift. The presence of an extra G is shown in comparison to the wild-type GFP sequence. **B)** Mutation analysis in biofilm by GFP* reversion assay. *M. smegmatis* strains harboring pFPVGFP* construct were seeded for biofilm formation with glass cover slip. Biofilm images from WT (left panel) and Δ *dnaE2* (right panel) strains were subjected to confocal microscopic analysis and fluorescent images were acquired.

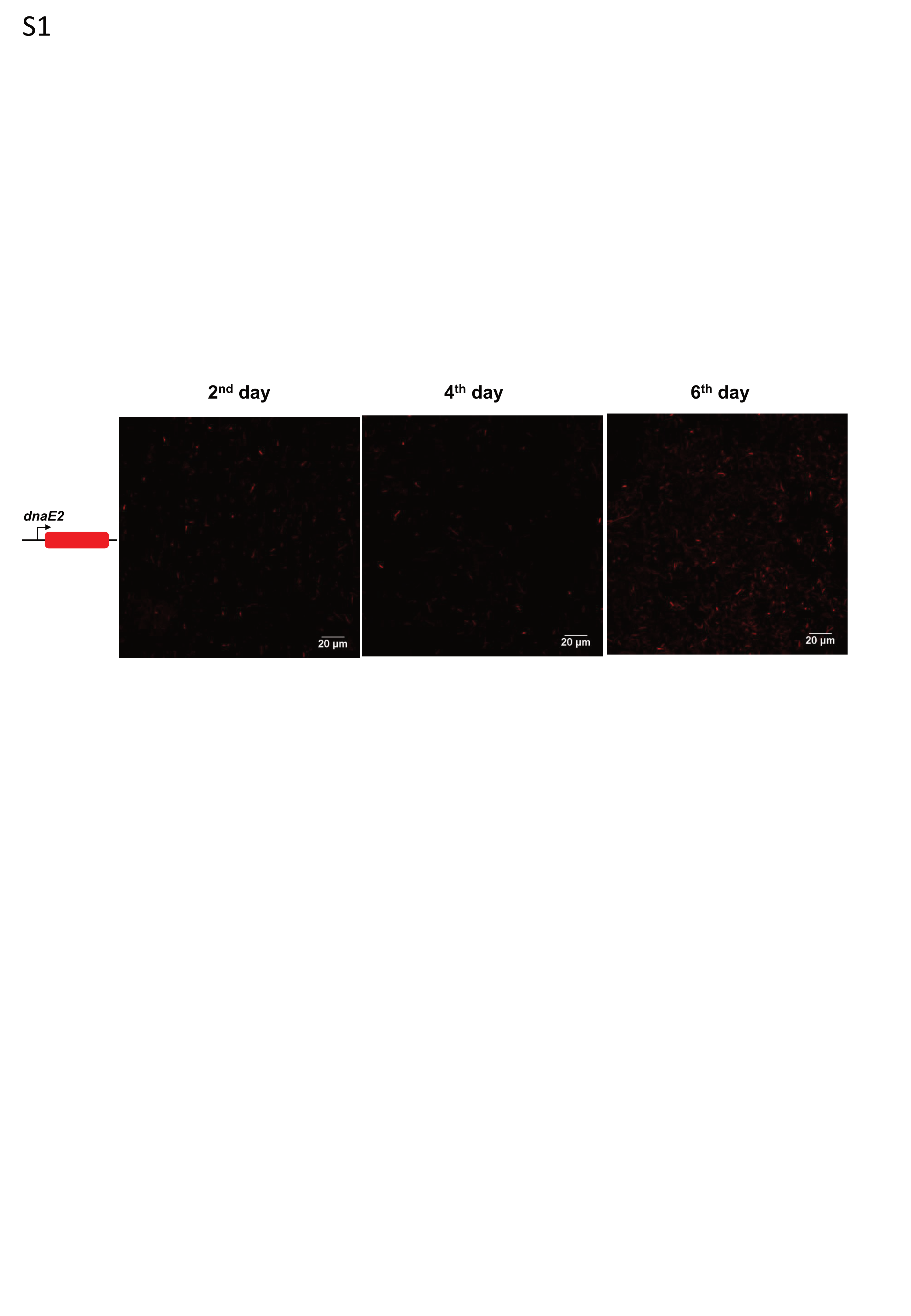

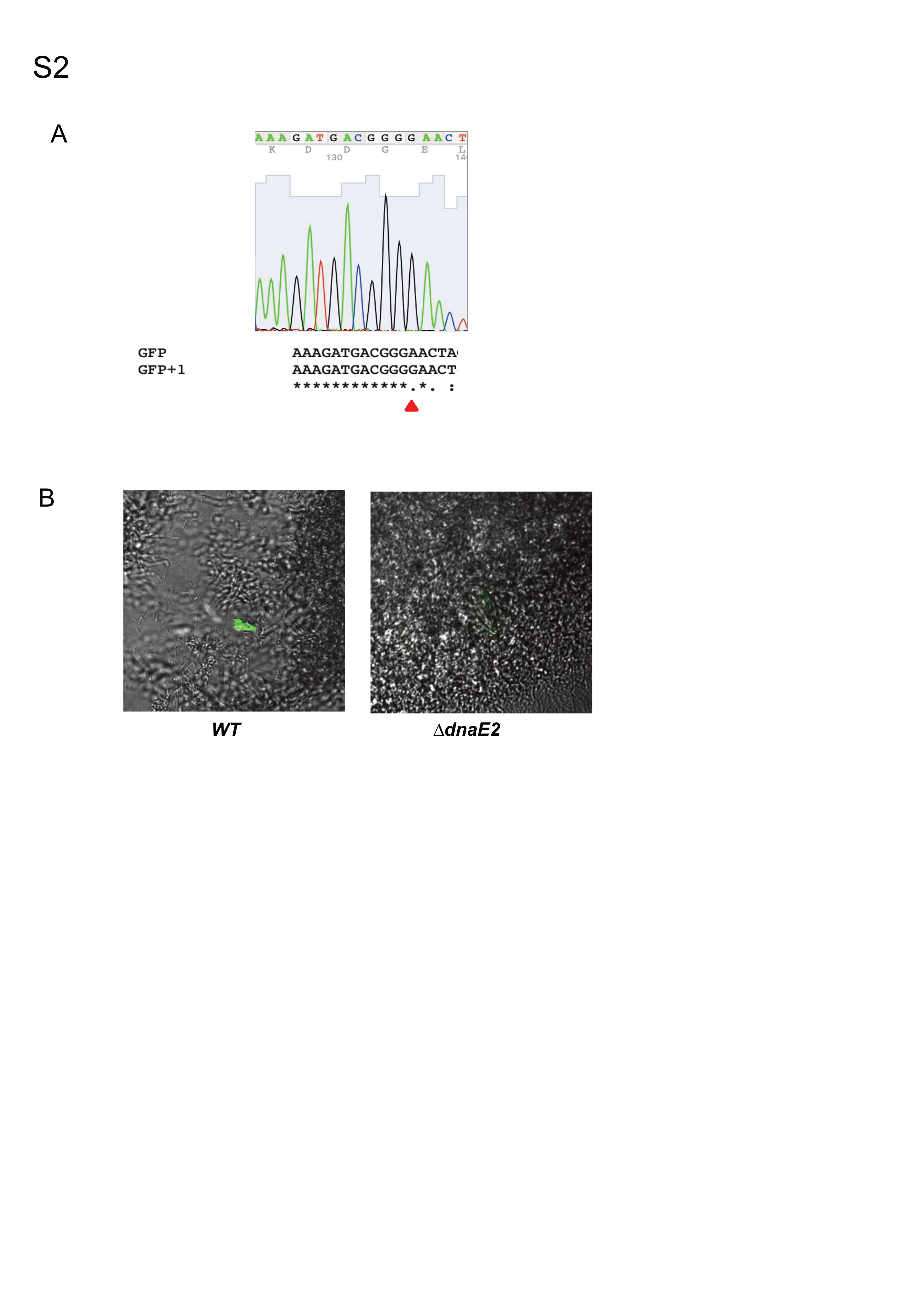
